# Discovery Stack Pilot Demonstrates the Feasibility and Outcomes of a Scientist-Designed Peer-Review Model That Separates Quality and Impact

**DOI:** 10.1101/2025.10.31.685758

**Authors:** Maureen A. McGargill, Beiyun C. Liu, Michael S. Kuhns, Daniel Mucida, Isabella Rauch, Lauren B. Rodda, Meghan A. Koch, Hugo Gonzalez, Ken Cadwell, Tanya S. Freedman, Tiffany C. Scharschmidt, Richard Sever, Jose Ordovas-Montanes, Andrew Oberst, Brooke Runnette, Matthew F. Krummel

## Abstract

Peer review serves as the cornerstone of scientific quality control. Yet, the current journal-centric system is hindered by long timelines, high publication costs, inconsistent review quality, systemic biases, and editorial gatekeeping. Notably, the system relies on misaligned measures of impact that are tethered to journal branding and conflate scientific rigor (*Quality*) with perceived significance (*Impact*). Here, we report findings from the Discovery Stack Pilot Study, which tested a scientist-designed, journal-independent peer review model. The Discovery Stack model integrates in-line reviewer comments to promote constructive feedback and separately evaluates scientific *Quality* and *Impact* using defined criteria. To examine feasibility and effectiveness, manuscripts were reviewed in parallel with traditional journal review. A total of 162 reviews were completed, and survey data from 86 participants were analyzed. The results showed that reviewers effectively evaluated *Quality* and *Impact* as separate dimensions, with *Quality* scores being more consistent across reviewers than *Impact* scores. Participants strongly supported the core elements of the Discovery Stack model and expressed enthusiasm for its broader adoption to enhance transparency, efficiency, and value in peer review. Future studies will explore integrating this model into a digital platform for reviewing and curating scientific discoveries to improve the production and dissemination of high-quality research.

## INTRODUCTION

Manuscript publication is the primary way researchers share new discoveries. Peer review prior to publication remains the central mechanism for evaluating the scientific rigor and validity of these findings (1). Researchers depend on this process to guide future studies, validate results, and uphold their professional reputation. Likewise, funding agencies, regulatory bodies, academic institutions, and industry stakeholders rely on peer-reviewed research, and the prestige of the journals in which it is published, to inform critical decisions on funding, policy, faculty promotion, media communication, product development, and patient care. In short, peer review underpins nearly every aspect of how scientific knowledge is generated, communicated, applied, and valued.

Despite its central role, the current journal-centric peer review and scientific publishing system often fails to meet the needs of scientists and society (2–6). A key limitation is the lack of a clear distinction between scientific *Quality* and *Impact*. *Quality* refers to the rigor and reproducibility of the data supporting a study’s conclusions, whereas *Impact* reflects the extent to which the findings advance understanding. When these dimensions are blurred, high-impact or hyped findings can overshadow weak evidence, while rigorous but incremental work is undervalued. Moreover, perceived *Impact* is often inferred from journal prestige rather than the intrinsic merit of the research itself (7, 8). Despite longstanding concerns regarding its validity, journal prestige is typically approximated using the Journal Impact Factor (JIF), a proprietary metric calculated by Clarivate (9–12). The JIF represents the mean number of citations received by articles published in a journal over a two-year period. However, citation distributions within journals are highly skewed, with a relatively small proportion of papers accounting for a large fraction of total citations. One analysis estimated that approximately 15% of articles generate 50% of citations, meaning that the journal average poorly reflects the influence of most individual publications (13). Furthermore, review articles or a single highly cited paper can substantially skew the JIF (12). Importantly, JIF may not be a reliable proxy for the methodological quality or reliability of individual publications. For example, higher JIFs are associated with higher retraction rates (11, 14). However, this pattern may partly reflect greater scrutiny and readership of articles published in highly visible journals, which increases the likelihood that errors are detected, rather than a difference in underlying research quality. Collectively, these dynamics distort how scientific contributions are valued, reinforcing journal reputation over scientific merit and undervaluing confirmatory studies that are essential for establishing confidence in foundational discoveries.

The traditional publishing process is also slow and inefficient. Manuscripts are considered by only one journal or journal family at a time, and each review cycle may take months. Across disciplines, the average time from submission to acceptance is approximately six months, but frequently extends well beyond a year (15, 16).

The quality, bias, and transparency of peer review are additional concerns. Reviews vary widely in depth and rigor, and critical flaws are sometimes missed (2–5, 16, 17). The lack of formal training and standardized guidelines contributes to this inconsistency (18). Reviewer anonymity, while intended to promote objectivity, can also shield bias and hostility from accountability. Together, these challenges undermine the effectiveness of peer review as a mechanism for quality control and diminish its value to authors.

Compounding these challenges, researchers perform peer review labor without compensation, while also paying to publish and access scientific literature. This model is inequitable, frustrating, and increasingly unsustainable, as publication numbers continue to rise without a proportional increase in the number of available reviewers (15, 19).

Although the growing prevalence of open-access preprint servers has improved accessibility to new findings (8, 20), these platforms often lack meaningful peer review or metrics of rigor. Consequently, readers face a new challenge, information overload, without effective mechanisms to evaluate, search, or filter studies based on *Quality* or *Impact*, making it difficult to determine which studies to trust and prioritize.

To address these challenges and accelerate scientific progress, the peer review process must be strengthened and modernized. We posit that an improved system should emphasize a “peer-improvement” mindset, positioning reviewers as collaborators focused on strengthening scientific rigor and benefiting the scientific community, rather than functioning as journal consultants determining binary publication eligibility. Such a system should apply standardized metrics to assess a study’s *Quality* and *Impact* as separate dimensions (21). Moreover, because *Impact* evolves over time through ongoing evaluation and influence on subsequent studies, this metric should remain dynamic and independent of journal branding.

The Discovery Stack Pilot Study tested the feasibility and effectiveness of a new peer review model built on these principles. The pilot evaluated the outcomes of separately assessing manuscript *Quality* and *Impact*, applying standardized metrics to independently measure and report both dimensions, and using an in-line commenting tool to facilitate constructive contextual feedback directly to authors and readers. The pilot also explored mechanisms for improving transparency, accountability, and timeliness, aiming to shorten the time from submission to dissemination. Findings from the Discovery Stack Pilot provide a foundation for further refinement and optimization of approaches to improve scientific publishing and peer review.

## RESULTS

### Discovery Stack Model

The Discovery Stack Pilot was designed to test a structured, multi-phase peer review process that independently evaluated scientific *Quality* from potential *Impact* using standardized assessment criteria. The process included three sequential phases: 1) *Quality* Review, 2) Author Response, and 3) *Impact* Review (**Fig 1A**). Each phase was built on the previous one, with *Impact* reviewers able to view compiled *Quality* review feedback and author responses. To ensure sufficient evaluations for each manuscript, reviewers who completed the *Quality* review phase also performed an independent assessment of the manuscript’s *Impact* using a separate evaluation form. Additional *Impact*-only reviewers were then recruited to increase the number and diversity of *Impact* assessments. Detailed procedures for each phase are provided in Materials and Methods.

**Fig 1.**
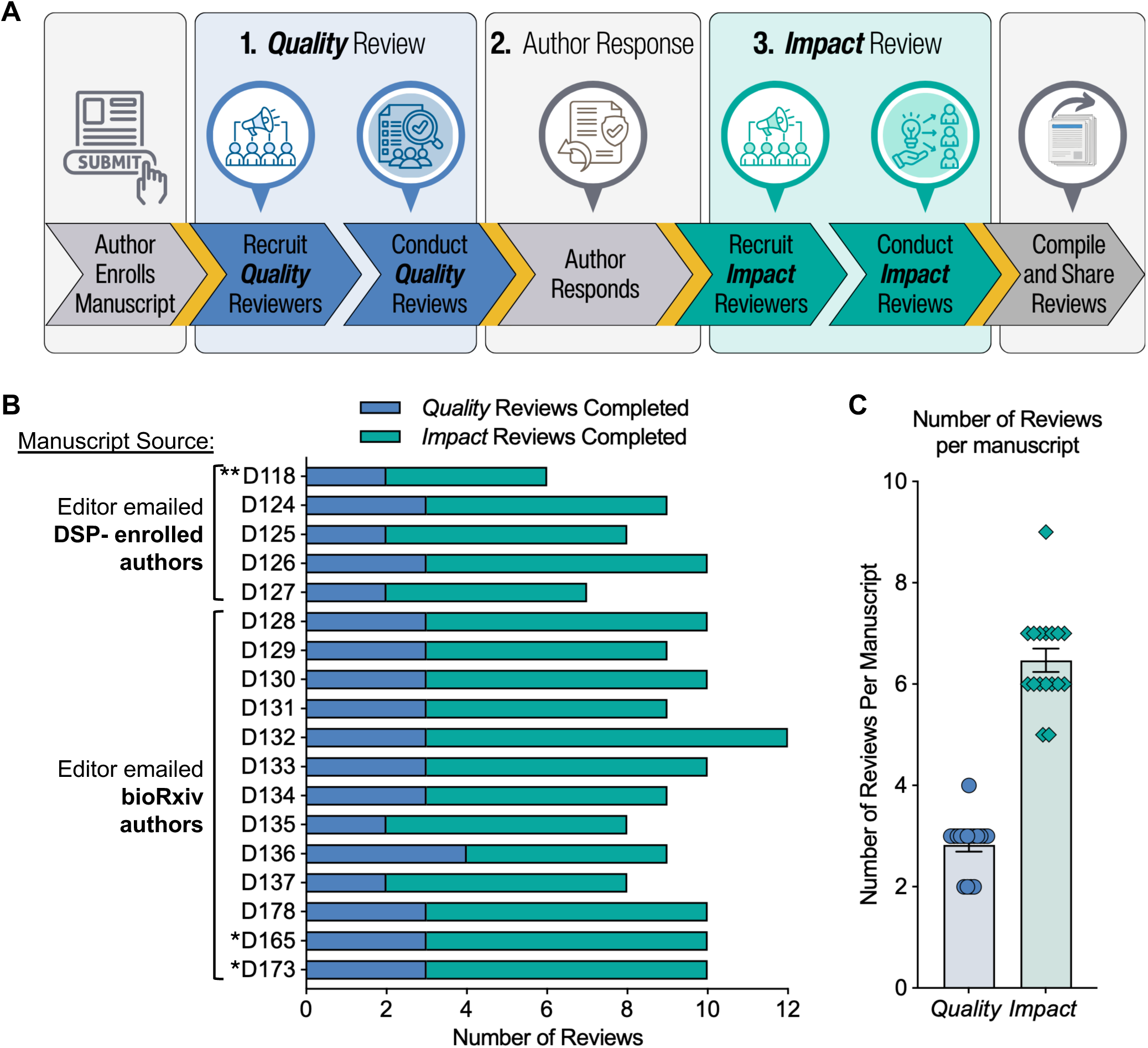
The Discovery Stack Model and Manuscript Enrollment. (**A**) The Discovery Stack model includes three sequential phases: 1) *Quality* Review, 2) Author Response, and 3) *Impact* Review. Image created in Canva. (**B**) Eighteen manuscripts were reviewed in the pilot. Manuscripts are grouped according to recruitment source and author participation, with the number of completed *Quality* and *Impact* reviews for each manuscript depicted. **Manuscript D118 was posted on SSRN rather than bioRxiv was excluded from some analyses due to compatibility issues between SSRN and Hypothes.is. *D165 and D173 were included in the pilot without author participation. (**C**) Number of *Quality* and *Impact* reviews completed per manuscript. The data underlying these figures are available in S1_Table and https://doi.org/10.5281/zenodo.2004122.

Manuscripts were eligible for enrollment if they: (1) were posted to a preprint server such as bioRxiv, which enabled testing of in-line comments through the Hypothes.is platform; (2) had been submitted to a traditional journal, allowing for comparison with conventional peer review; and 3) were in the fields of immunology or cancer biology, enabling us to leverage the subject expertise of the scientific advisory board and participating reviewers.

To identify eligible manuscripts, initial invitations were sent to 116 individuals who had previously registered to participate as authors or reviewers in the Discovery Stack Pilot, which resulted in five enrolled manuscripts. Outreach was then expanded to corresponding authors of bioRxiv preprints that met the above criteria. The most recent preprints were prioritized to maximize the likelihood that manuscripts had not yet been accepted for journal publication. Using these criteria, 118 bioRxiv authors were contacted, resulting in 11 more enrolled manuscripts. Additionally, two bioRxiv preprints for which no author response was received were included in the pilot, as reviewers with appropriately matched expertise had already been identified and agreed to participate.

In total, 18 manuscripts were enrolled (**Fig 1B**), generating 162 reviews: 50 *Quality* and 112 *Impact* reviews (**Fig 1B-C**). Each manuscript received at least two *Quality* and five *Impact* reviews, with an average of 2.8 *Quality* and 6.5 *Impact* reviews per manuscript. Because we hypothesized that *Impact* assessments would be inherently more subjective and variable, we aimed to recruit three *Quality* reviewers and six *Impact* reviewers per manuscript. Most manuscripts met or exceeded this goal, demonstrating the feasibility of enrolling manuscripts, recruiting reviewers, and completing both *Quality* and *Impact* assessments.

The Discovery Stack Pilot was designed as a feasibility study to evaluate the implementation of this model under real-world conditions. The primary objectives were to assess recruitment, workflow execution, and whether the model could generate actionable insights into key aspects of peer review, including the separation of *Quality* and *Impact* and participant perceptions of standardized metrics. As such, results should be interpreted in the context of a pilot study. In addition, because enrollment relied on voluntary participation from both previously enrolled Discovery Stack participants and authors of recent bioRxiv preprints, the study population may have been enriched for individuals more receptive to alternative peer review approaches. This, together with the relatively small cohort size, should be considered when interpreting perception-based outcomes. Larger, more systematically recruited cohorts will be required to confirm and extend these findings.

### Reviewers Successfully Distinguished *Quality* from *Impact*

A central feature of the Discovery Stack model is the independent evaluation of scientific *Quality* and *Impact* using structured, standardized assessment criteria. To implement this framework, reviewers completed assessment forms evaluating defined attributes of scientific *Quality* and *Impact.* The *Quality* Assessment Form consisted of six short-answer questions and 13 Likert-scale items addressing experimental design, controls, statistical analysis, reproducibility, and whether the data supported the stated conclusions (**Fig 2**). The *Impact* Assessment Form included a short-answer question, a multiple-choice question identifying features contributing to a manuscript’s *Impact*, and five Likert-scale items evaluating transformative potential, generalizability, technological advancement, and mechanistic insight (**Fig 3**). To generate quantitative metrics for filtering and prioritizing manuscripts by *Quality* and *Impact*, composite scores were generated from reviewer responses. Each question was weighted according to its relative importance to either the *Quality* or *Impact* of a manuscript (**Fig S1A-B**). Weighted responses from each reviewer were averaged to yield a single composite *Quality* and a single composite *Impact* score per reviewer per manuscript. Composite scores across reviewers were then averaged to produce a final *Quality* and *Impact* score for each manuscript (**S1 Table**; see Materials and Methods for full scoring details).

**Fig 2.**
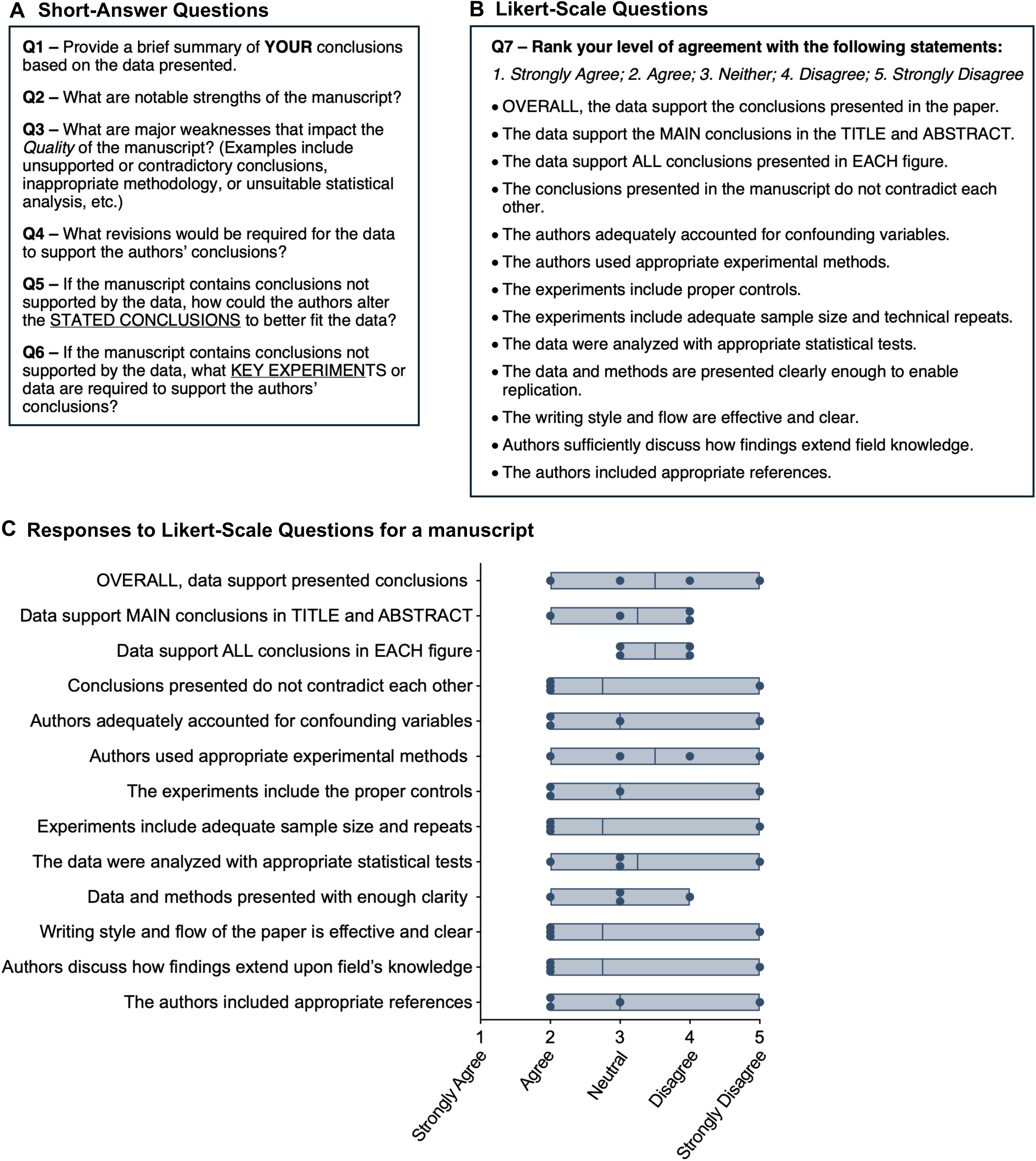
*Quality* Review Assessment Form. The *Quality* Review Assessment Form included (A) six short-answer questions, followed by (B) a set of Likert-scale items designed to evaluate specific attributes regarding the rigor and reproducibility of the data. (C) Scores from all reviewers were compiled, graphed, and shared with the authors.

**Fig 3.**
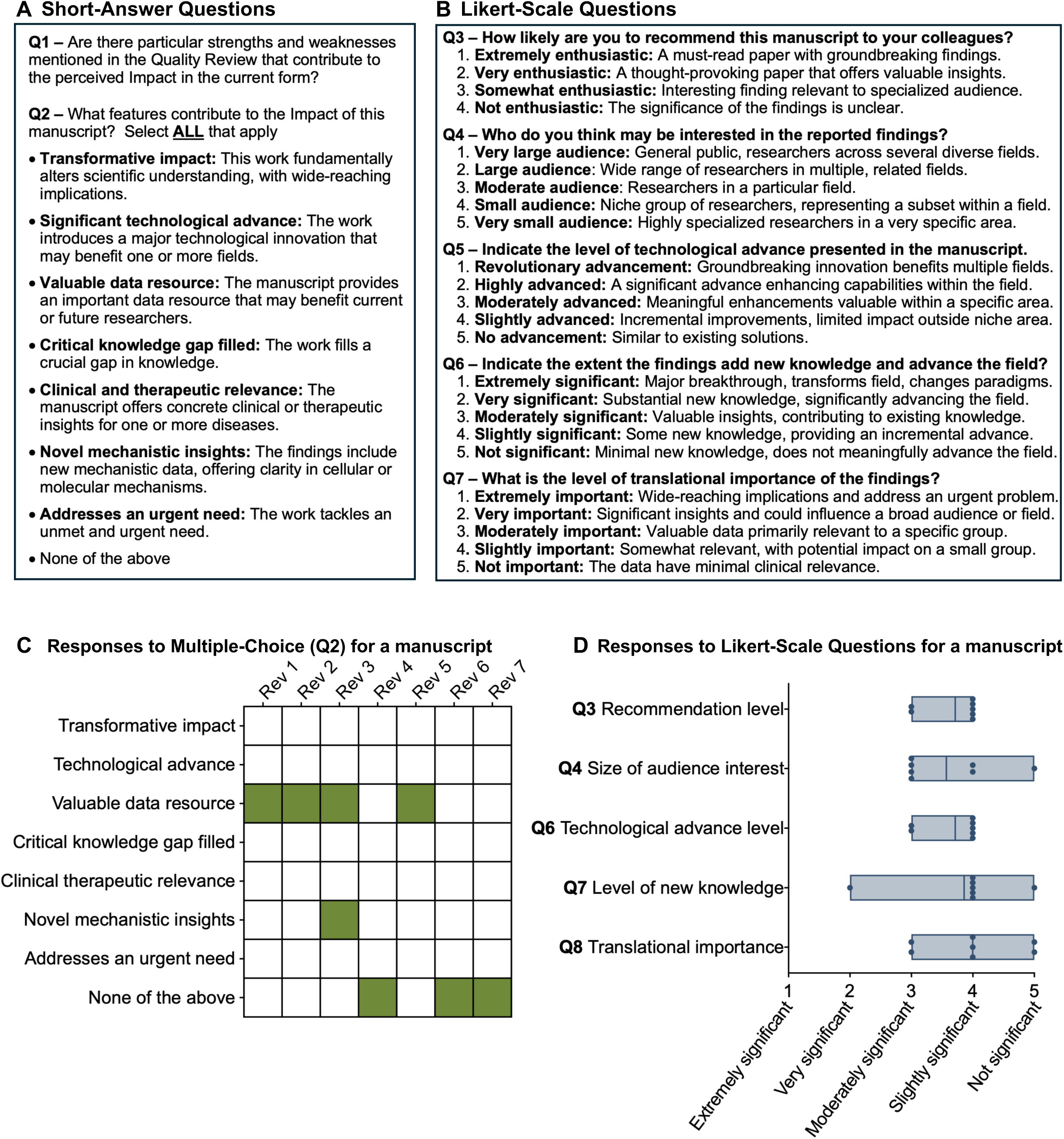
*Impact* Review Assessment Form. (A) The *Impact* Review Assessment Form included: a short-answer question asking whether specific strengths or weaknesses identified in the *Quality* Review influenced their perception of the manuscript’s *Impact*, a multiple-choice question allowing reviewers to select features they believed contributed to the manuscript’s *Impact*, and (B) five Likert-scale questions assessing key dimensions of the manuscript’s *Impact*. (C) Multiple choice and (D) Likert-scale responses were compiled, graphed, and shared with the authors.

To determine whether reviewers evaluated *Quality* and *Impact* as distinct dimensions, we examined the relationship between composite *Quality* and *Impact* across manuscripts. First, composite *Quality* and *Impact* scores for each manuscript were plotted according to the JIF of the journal to which each manuscript was published or submitted, which served as a proxy for the author’s perception of *Impact* (**Fig 4A**). This visualization showed several manuscripts rated high in *Quality* but low in *Impact*, suggesting that reviewers were able to uncouple their assessment of *Impact* from *Quality*. To formally test this, we quantified the relationship between these two dimensions. Because each manuscript received more *Impact* than *Quality* reviews, the primary analysis was restricted to reviewers who provided both *Quality* and *Impact* scores for the same manuscript to minimize differences driven by unequal sample sizes and reviewer variability. Pearson’s correlation showed a significant positive association between *Quality* and *Impact* (r = 0.73, *p*=0.0006), and linear regression confirmed that significantly predicted *Impact* (**Fig 4B**; β = 0.70, R² = 0.53 *p*=0.0006). As expected, poor-*Quality* manuscripts are unlikely to be considered impactful. However, residual analysis indicated substantial divergence. Ten of 18 manuscripts (56%) had *Impact* scores outside the 95% confidence interval (**Fig 4B**), with residuals ranging from −0.99 to +0.53 *Impact* points (**Fig 4C**). The standard deviation of the residuals (SD = 0.41) further demonstrated variation around the regression line, indicating that reviewers’ *Impact* evaluations frequently diverged from predictions based solely on *Quality*. Analyses including all reviewer scores yielded comparable results (r = 0.78, *p*=0.0001; β = 0.68, R² = 0.60, *p*=0.0001).

**Fig 4.**
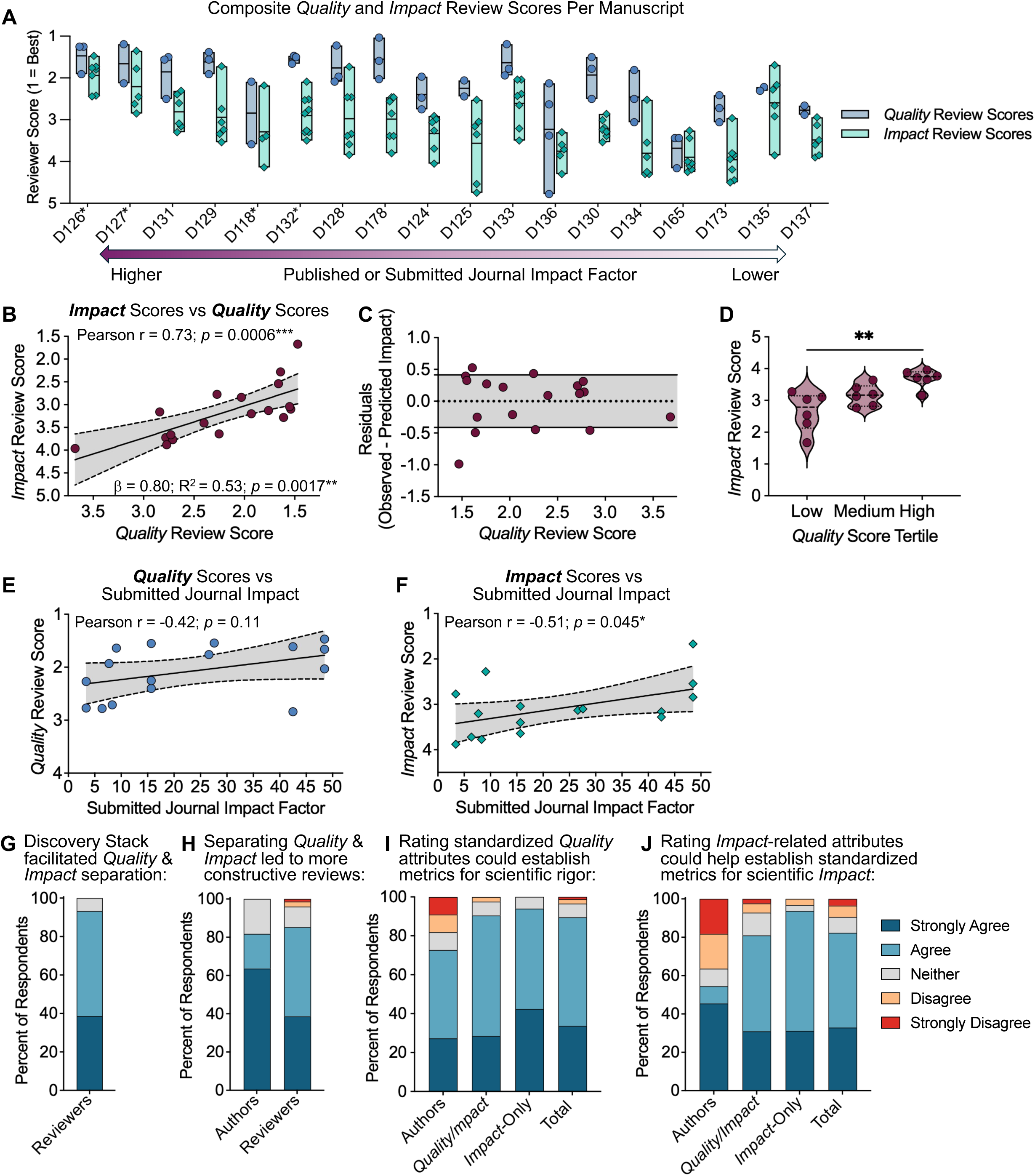
Effective Separation of *Quality* and *Impact* with Strong Support for Standardized Metrics. (**A**) Composite *Quality* and *Impact* scores arranged by impact factor of the journal the manuscript was published in. *Manuscripts that were not published in a journal at the time of the final analysis are denoted by an asterisk. (**B**) The relationship between composite *Quality* and *Impact* scores for each manuscript was tested using Pearson’s correlation and linear regression, restricting analysis to reviewers who completed both assessments. The shaded region shows the 95% confidence interval (CI) of the regression line. (**C**) Residuals from the linear regression model to assess variation in *Impact* not explained by *Quality*. Shaded region represents ±1 standard deviation (SD = 0.41). (**D**) Manuscripts were grouped into tertiles based on *Quality* scores and *Impact* scores were compared across tertiles using Kruskal–Wallis tests (H = 10.4, *p*=0.0043), followed by Dunn’s multiple comparisons. (**E-F**) Pearson’s correlation was used to evaluate the relationship between *Quality* (**E**) or *Impact* (**F**) scores and the JIF of the journal to which each manuscript was submitted. The submission journal of D165 and D173 was unknown, excluding them from the analysis. (**G-J**) Survey data from *Quality/Impact* reviewers (n=42), *Impact*-only reviewers (n=33), and authors (n=11) were tallied, and the percentage of each group selecting each response is shown. The data underlying these figures are available in S1_Table and https://doi.org/10.5281/zenodo.2004122.

To further assess the independence between *Impact* and *Quality*, manuscripts were grouped into tertiles by ranked composite *Quality* scores, and mean *Impact* scores were compared the across groups. Mean *Impact* scores increased with higher *Quality*, but significant differences were observed only between the lowest and highest tertiles, with substantial overlap between adjacent groups (**Fig 4D**). The residual variation (SD = 0.41) was nearly as large as the average difference in *Impact* between tertiles (0.51–0.53), indicating that manuscripts with comparable *Quality* scores frequently differed in their *Impact* ratings.

Next, we examined whether reviewer scores aligned with the JIFs to which the manuscripts were submitted. It is important to note that Discovery Stack reviewers were unaware of which journals the authors selected. *Quality* scores did not significantly correlate with JIFs (**Fig 4E**; r = −0.42, *p*= 0.11), whereas *Impact* scores were significantly correlated (**Fig 4F**; r = −0.51, *p*=0.045). Again, analyses including all reviewer scores yielded comparable results (*Quality* r = −0.44, *p*= 0.092; *Impact* r = −0.57, *p*= 0.021). These findings suggest that reviewers’ perceptions of *Impact* aligned more closely with the authors’ expectations of significance than their *Quality* assessments. Although the modest sample size may have limited our ability to detect a statistically significant association between *Quality* scores and JIFs, these findings are consistent with evidence that JIF is a poor surrogate for scientific reliability (11). Similarly, when manuscripts were grouped into JIF tiers based on the submitted or published journal, manuscripts receiving better *Impact* scores generally aligned with higher JIF tiers, whereas *Quality* scores showed weaker correspondence (**Fig S2**). Together, these results demonstrate that reviewers distinguished between *Quality* and *Impact* as separate but complementary dimensions of manuscript evaluation.

A limitation of this analysis is that comparisons between Discovery Stack scores of revised manuscripts to JIFs in which manuscripts were ultimately published were not possible, as reviewers only assessed initial submissions. In addition, at the time of the final analysis, only 14 of 18 manuscripts were accepted for journal publication, which further limited comparisons between Discovery Stack scores and the JIF of the journals that ultimately accepted the manuscripts. Additionally, it is possible that reviewers inferred the tier of journal selected by authors based on formatting of the preprint.

### Widespread Endorsement for Standardized Metrics and Separate Evaluation

To evaluate participant perceptions following completion of the pilot study, reviewers and authors completed surveys assessing key features of the Discovery Stack model. Since separating *Quality* from *Impact* was a central component of the framework, we examined whether participants believed the Discovery Stack model effectively supported this distinction and whether doing so enhanced the review process. The vast majority of reviewers (93%) agreed or strongly agreed that the model effectively supported separate assessments of *Quality* and *Impact* (**Fig 4G**). Moreover, 85% of all participants agreed that separating these dimensions led to more constructive and insightful evaluations than traditional reviews (**Fig 4H**).

We also examined perceptions of rating standardized attributes as a means to establish quantitative metrics for scientific rigor and *Impact*. Support for standardized metrics was compelling, with 90% of participants agreeing that standardized *Quality* ratings could generate meaningful metrics of rigor (**Fig 4I**), and 82% agreeing that standardized *Impact* ratings could capture perceived significance (**Fig 4J**). Reviewer support exceeded author support for both metrics (92% vs. 72% for *Quality*; 86% vs. 55% for *Impact*), though the author sample was smaller than the reviewer sample (n=11 vs. n=75). These findings demonstrate broad endorsement of two core elements of the Discovery Stack model: (1) the separation of *Quality* and *Impact,* and (2) the use of standardized metrics to increase transparency, improve review *Quality*, and reduce reliance on journal branding as a measure of scientific value. An important consideration is that participant support for standardized metrics may also reflect endorsement of the structured assessment framework itself, including the use of clearly defined evaluation criteria to guide reviewer assessments of *Quality* and *Impact*. Such guidance is uncommon in traditional peer review and may have contributed to perceptions of improved clarity, consistency, and transparency.

Because participation in this pilot was voluntary, and recruitment relied in part on prior engagement and professional networks, we considered the possibility that responses may have been influenced by prior familiarity with the Discovery Stack model. To assess this, survey responses from participants new to the study (NEW) were compared to those from participants previously enrolled in the pilot (DSP-enrolled). Across survey questions, responses were highly similar between new and previously enrolled participants, with comparable median scores and uniformly small effect sizes (Table in **S2 Table**). After adjustment for multiple comparisons, only one question showed a statistically significant difference in responses, while all others were not significant. Visualization of mean differences with bootstrap confidence intervals further demonstrated that most comparisons were centered near zero with overlapping confidence intervals, indicating minimal differences between groups (**Fig S3A-B)**. Together, these results suggest that prior familiarity with the Discovery Stack model did not meaningfully influence participant responses. Nevertheless, because participation in the study was voluntary, participants may be more receptive to alternative models of peer review than the broader scientific community, which should be considered when interpreting these results.

### *Impact* Reviews Exhibit Greater Variability Than *Quality* Reviews

Visual inspection of side-by-side *Quality* and *Impact* scores suggested that, within individual manuscripts, *Impact* scores varied more than *Quality* scores (**Fig 4A**). Therefore, we compared reviewer score dispersion using three complementary metrics: standard deviation (SD), range (maximum - minimum score), and interquartile range (IQR; difference between the 75th and 25th percentile). Across all three metrics, *Impact* scores consistently exhibited greater dispersion than *Quality* scores (**Fig 5A-C**). To account for unequal numbers of *Impact* and *Quality* reviews per manuscript, we randomly subsampled the *Impact* scores to match the number of *Quality* scores for each manuscript and recalculated the variability metrics across 5,000 iterations. Bootstrap distributions of the differences in variability between *Impact* and *Quality* scores were shifted above zero across all three metrics, consistent with greater variability in *Impact* assessments even after subsampling *Impact* reviews to match the number of *Quality* reviews per manuscript (**Fig 5D**; Table in **S3_Table**). These findings underscore the value of evaluating *Quality* and *Impact* as distinct dimensions and support the need for a greater number of *Impact* reviewers to capture the broader range of perspectives on scientific significance.

**Fig 5.**
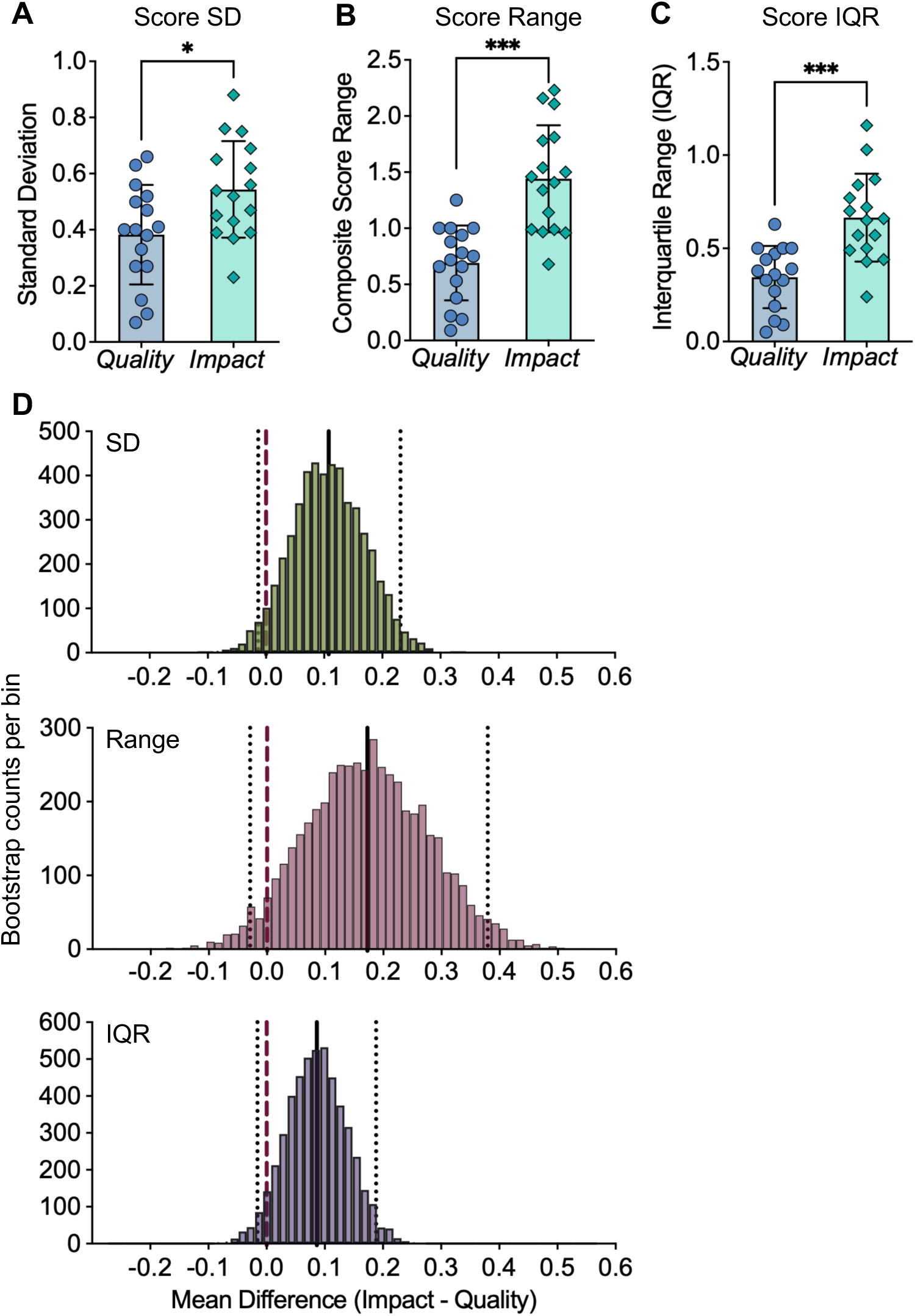
*Impact* Reviews Exhibit Greater Variability Than *Quality* Reviews. Variability between composite *Quality* and *Impact* scores, within each manuscript, was quantified using (**A**) standard deviation (SD), (**B**) range, and (**C**) interquartile range (IQR), and compared using Wilcoxon matched-pairs signed-rank test. *p*<0.05*; *p*<0.001***. (**D**) A subsampling approach was performed to compare variability between *Impact* and *Quality* scores. For each manuscript, *Impact* scores were randomly subsampled to match the number of *Quality* scores, and variability metrics were recalculated for each subsampled dataset. The subsampling procedure was repeated for 5,000 iterations to generate empirical distributions of the mean differences (*Impact* - *Quality*) for each variability metric. Confidence intervals and one-sided p-values were derived from these distributions, with p-values defined as the proportion of iterations in which the mean difference (*Impact* - *Quality*) was less than or equal to zero (Table in **S3_Table**). Manuscript D136 was excluded as an outlier (z-score = −3.14, >3 SD from mean difference) and D118 was excluded as additional *Impact* reviewers were not recruited due to Hypothes.is compatibility issues, resulting in a final sample size of n=16 manuscripts for panels A-D. The data underlying these figures are available in S3_Table and https://doi.org/10.5281/zenodo.2004122.

### Identity Disclosure May Be Associated with Greater Perceived Transparency and Higher *Impact* Scores

To promote transparency, reviewers were encouraged to disclose their identity to authors and co-reviewers, although anonymity remained an option to preserve the integrity of the review process. Approximately half of reviewers disclosed their identity: 50% of *Quality* reviewers and 54% of *Impact* reviewers (**Fig S4A**). Interestingly, a greater proportion of trainees (73%) than principal investigators (47%) identified themselves (**Fig S4B**). The proportion of identified reviewers varied substantially across manuscripts (**Fig S4C**; range: 0-100%), suggesting that factors such as authorship or perceived study *Quality* may have influenced identity disclosure decisions. The most common reasons for remaining anonymous were familiarity with the authors and concern about professional repercussions (**Fig S4D**).

Among the 11 authors who responded, eight (73%) agreed that the Discovery Stack review was more transparent than traditional review (**Fig S4E**). Authors whose manuscripts had a higher proportion of identified reviewers tended to perceive greater transparency (**Fig S4F**). Given the limited number of author responses, these findings should be considered preliminary observations rather than definitive evidence.

*Quality* reviewers’ feedback was shared with authors and *Impact*-only reviewers, allowing both groups to evaluate whether reviewer identity influenced scoring. Most *Impact*-only reviewers (89%) reported observing no clear differences in scores between identified and anonymous reviewers (**Fig S4G**). The small number of author responses were mixed, with six of 10 reporting no clear difference between anonymous and identified reviewers, while the remaining four stated that identified reviews were more constructive than anonymous reviews. These mixed responses warrant further study in larger cohorts.

To directly evaluate whether reviewer identity influenced scores, we compared individual scores from anonymous and identified reviewers across all manuscripts (unpaired) and within manuscripts (paired). The paired analysis assessed whether, for a given manuscript, scores differed between anonymous and identified reviewers. There were no significant differences in *Quality* scores between the two groups (**Fig S4I-J**), but *Impact* scores were significantly higher among identified reviewers (**Fig S4K-L**). These findings were consistent in the unpaired analysis across all manuscripts and within manuscripts in the paired analysis. Additionally, the higher *Impact* scores from identified reviewers was not explained by reviewer career stage (**Fig S4M**).

In summary, identity disclosure was associated with higher *Impact* scores and may influence perceptions of transparency. Given the small number of author responses, these observations should be considered preliminary. Nevertheless, they raise the possibility that reviewers may be more likely to identify themselves when giving favorable evaluations, or alternatively, that agreeing to disclose identity may incline reviewers toward softer *Impact* assessments. These observations warrant further investigation in larger studies of how identity disclosure influences both the review process and its perception.

### Discovery Stack Model Delivers a Better Experience than Traditional Review

A central goal of the Discovery Stack Pilot was to assess whether participants believed that the model offered a better experience than traditional peer review. Overall, a strong majority (83%) rated their experience as “*much better”* or “*slightly better”*, while only 3.5% rated it as worse (**Fig 6A**). Positive ratings were highest among *Impact*-only reviewers (94%), followed by authors (82%; 9 of 11), and *Quality/Impact* reviewers (74%). It is important to consider that unlike traditional peer review, the Discover Stack Pilot did not involve editorial accept or reject decisions, which may contribute to the more favorable author perceptions of the review experience. The lower satisfaction among *Quality/Impact* reviewers may reflect the additional effort required to complete both review phases and learn the Hypothes.is platform.

**Fig 6.**
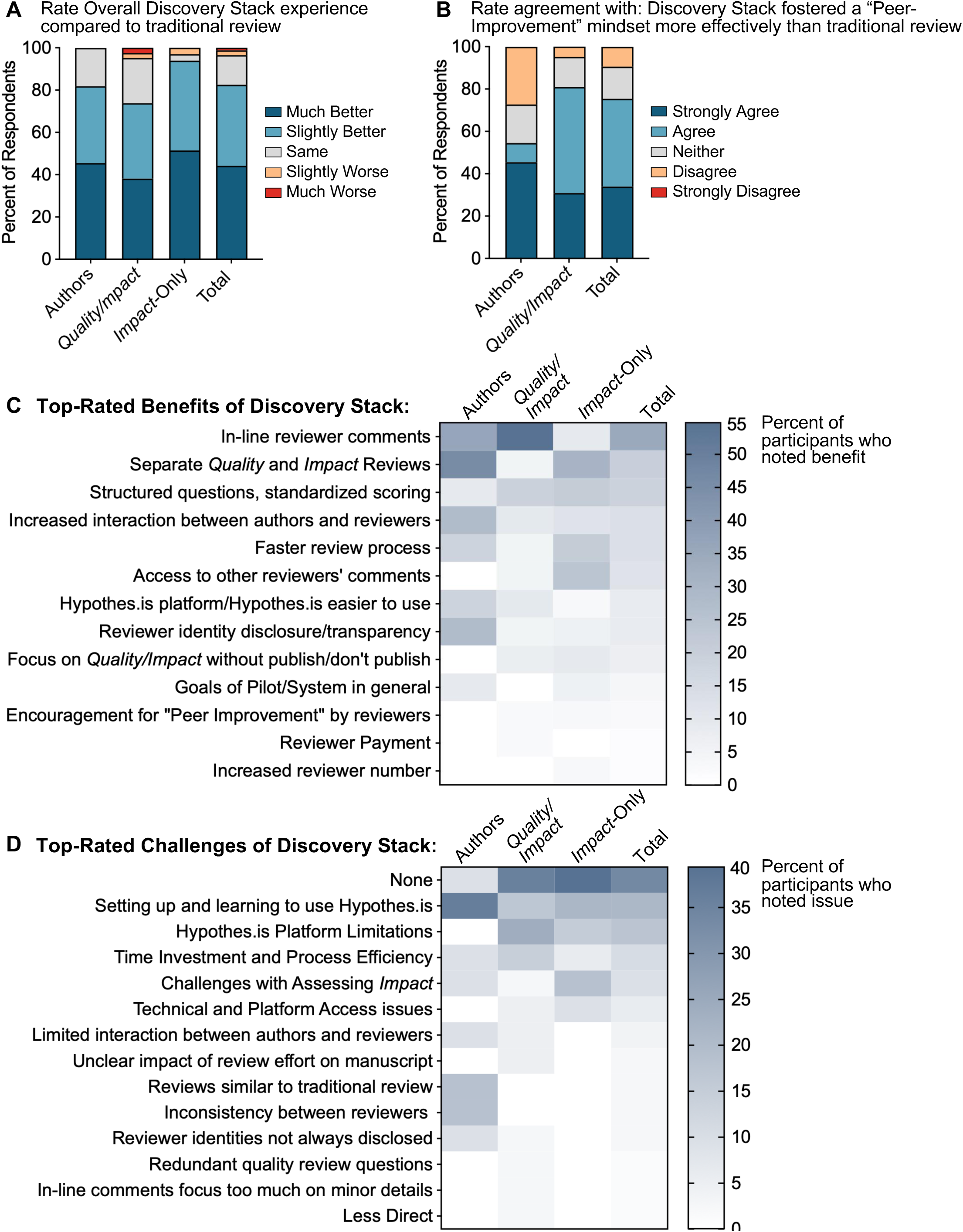
High Satisfaction with the Discovery Stack Model. (**A**) Participants rated their overall experience with the Discovery Stack model compared to traditional review. The percent of authors (n=11), *Quality/Impact* reviewers (n=42), and *Impac*t-only reviewers (n=33) selecting each rating is shown. (**B**) Authors (n=11) and *Quality/Impact* reviewers (n=42) were asked whether the Discovery Stack model fostered a greater “Peer-Improvement” mindset than traditional peer review. (**C**) Participants were asked in an open-ended question what aspects of the Discovery Stack model they found most beneficial compared to traditional review. A total of 74 responses were collected, categorized into key thematic areas, and the percent of participants (n=86) who mentioned each theme is shown. All responses are shown. (**D**) Participants identified aspects of the model they found most challenging. A total of 56 open-ended responses were collected, categorized into thematic categories, and the percentage of participants (n=86) mentioning each challenge is shown. All responses are shown. The data underlying these figures are available at https://doi.org/10.5281/zenodo.2004122.

Only three participants reported a worse experience, citing the need to consult setup instructions, perceived platform complexity, or uncertainty about how their reviews affected manuscript outcomes. These challenges are typical of new systems and are expected to diminish with familiarity. Additionally, because the pilot ran in parallel to traditional review, authors were not required to respond to Discovery Stack feedback, which limited reviewers’ insight into the impact of their efforts.

A core tenet of the Discovery Stack model is to promote a “peer-improvement” mindset, encouraging reviewers to provide constructive, actionable feedback that enhances scientific rigor. This principle was operationalized through multiple components of the model, including reviewer onboarding that explicitly emphasized the value of constructive, peer-improvement-oriented feedback, as well as the structured *Quality* and *Impact* assessment forms, in-line annotation, and the separation of *Quality* and *Impact* review phases. Most participants (75%) agreed that the Discovery Stack model fostered this mindset more effectively than traditional reviews, with stronger agreement among reviewers (81%) than authors (55%; 6 of 11) (**Fig 6B**). Because the survey assessed perceptions of the model as a whole, this result likely reflects the combined contributions of these components, including the reviewer briefing.

To gain qualitative insight, participants were asked open-ended questions about the most beneficial and most challenging aspects of the Discovery Stack model compared to traditional review. Consistent with the positive ratings, more participants cited benefits (n=74) than challenges (n=56). The most common benefits were in-line commenting (35%), separation of *Quality* and *Impact* reviews (20%), and standardized questions and scoring (18.6%) (**Fig 6C**). The most frequent challenges were setting up and learning Hypothes.is (20%), Hypothes.is limitations (9%), and time demands (11%) (**Fig 6D**).

Although challenges with the Hypothes.is tool were most frequently cited, in-line commenting was also the most reported benefit, highlighting both its value as a core feature and the need for technical enhancements. With 82% of participants reporting a better experience than traditional review, these findings provide strong support for Discovery Stack and its potential for broader adoption with continued optimization.

### In-line Commenting Improves Review Clarity, Collegiality, and Efficiency

Given that in-line commenting was both the most cited benefit and key area for improvement, we evaluated its effectiveness in improving review clarity, collegiality, and efficiency. The Hypothes.is tool was selected because it enabled contextual annotation of bioRxiv-hosted preprints within private groups, allowing reviewers to leave feedback directly on the manuscript text. Responses from both reviewers and authors strongly supported this feature. Most authors and *Impact*-only reviewers (71%) agreed that in-line comments were more collegial and constructive than traditional reviews (**Fig 7A**). Notably, all authors that responded (10 of 10) found in-line comments easier to respond to and more helpful for identifying needed revisions than traditional review summaries (**Fig 7B**).

**Fig 7.**
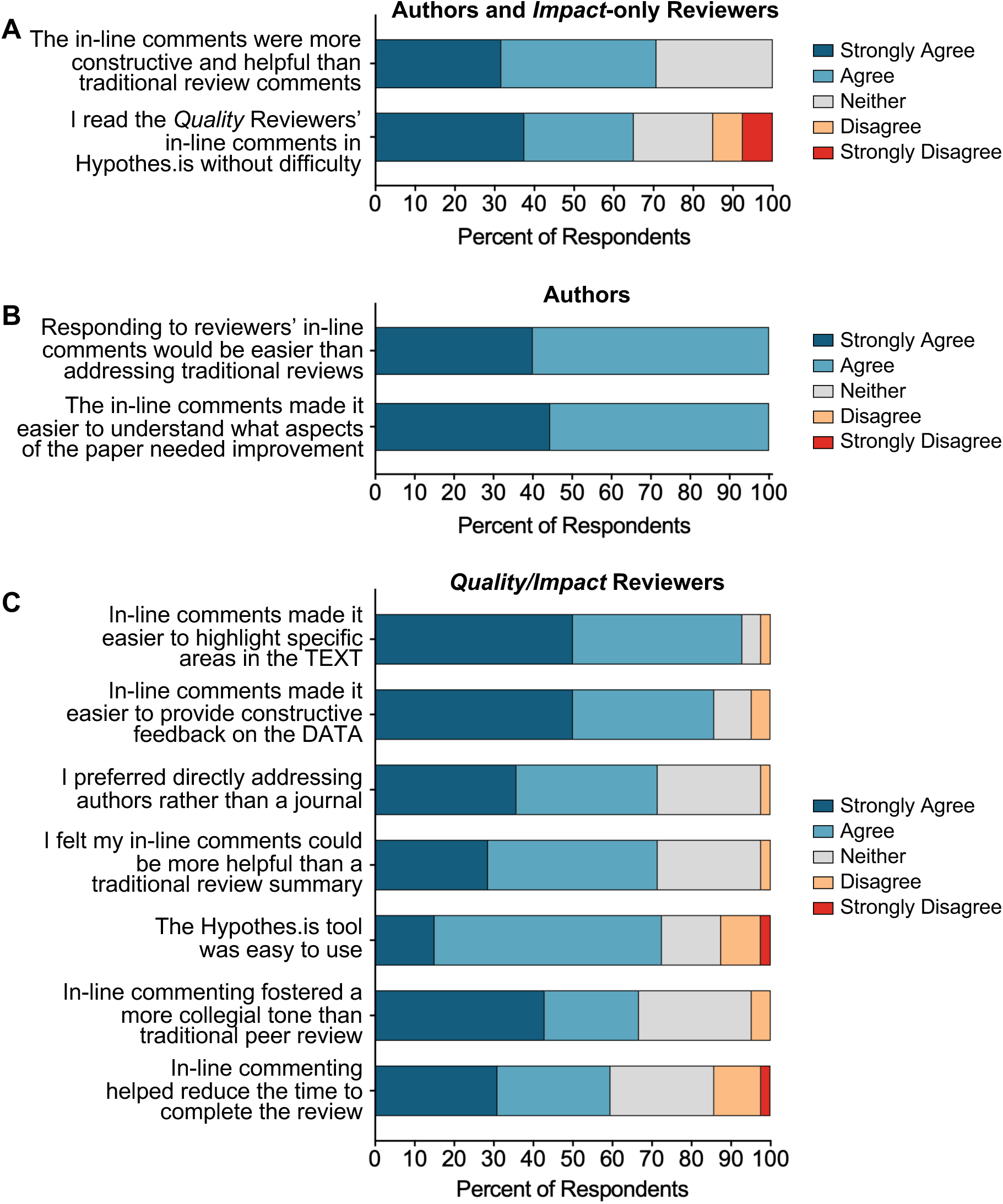
In-line Commenting Improves Review Clarity, Collegiality, and Efficiency. (**A**) Authors (n=8-9) and *Impact*-only reviewers (n=32) were asked whether they were able to read the *Quality* reviewers’ in-line Hypothes.is comments without difficulty, and whether they found the comments more constructive and helpful than those typically received through traditional review. (**B**) Authors (n=8-9) were asked whether responding to reviewers’ in-line comments would be easier than addressing traditional review comments, and whether the in-line format helped clarify which aspects of the paper needed improvement. (**C**) *Quality/Impact* reviewers (n=40-42) were asked a series of questions focused on providing in-line comments using the Hypothes.is platform. The data underlying these figures are available at https://doi.org/10.5281/zenodo.2004122.

Among *Quality/Impact* reviewers, 93% agreed that in-line comments made it easier to highlight errors in logic or clarity, 86% felt in-line commenting improved the constructiveness of feedback, and 71% preferred addressing comments directly to the authors, believing that their in-line comments would be more helpful than a traditional review summary (**Fig 7C**). Although the Hypothes.is tool was new to most reviewers, 73% reported that it was easy to use, and 60% agreed that in-line commenting improved review efficiency. Additionally, 67% agreed that in-line commenting helped them adopt a more collegial tone (**Fig 7C**). Together, these findings indicate that in-line commenting is both feasible and effective, providing clearer, more constructive, and more collegial feedback than traditional review.

### Authors Find Discovery Stack *Quality* Reviews Rigorous and Helpful

We next evaluated whether the Discovery Stack model effectively directed reviewers to focus on scientific rigor by asking authors about their perception of the *Quality* reviewer feedback. Most authors (82%; 9 of 11) agreed that *Quality* reviews focused on scientific rigor, and 73% (8 of 11) found the standardized *Quality* and *Impact* scores helpful for understanding how their manuscript was evaluated (**Fig 8A**). Likewise, 82% (9 of 11) agreed the *Quality* Assessment Form effectively summarized the key strengths and weaknesses, and more than 60% (5 of 8) reported revising their manuscript based on reviewers’ feedback. In-line commenting was also well received, with seven of eight (88%) authors agreeing that the opportunity to interact with reviewers through Hypothes.is provided a good method to clarify feedback and expedite the review process (**Fig 8A**).

**Fig 8.**
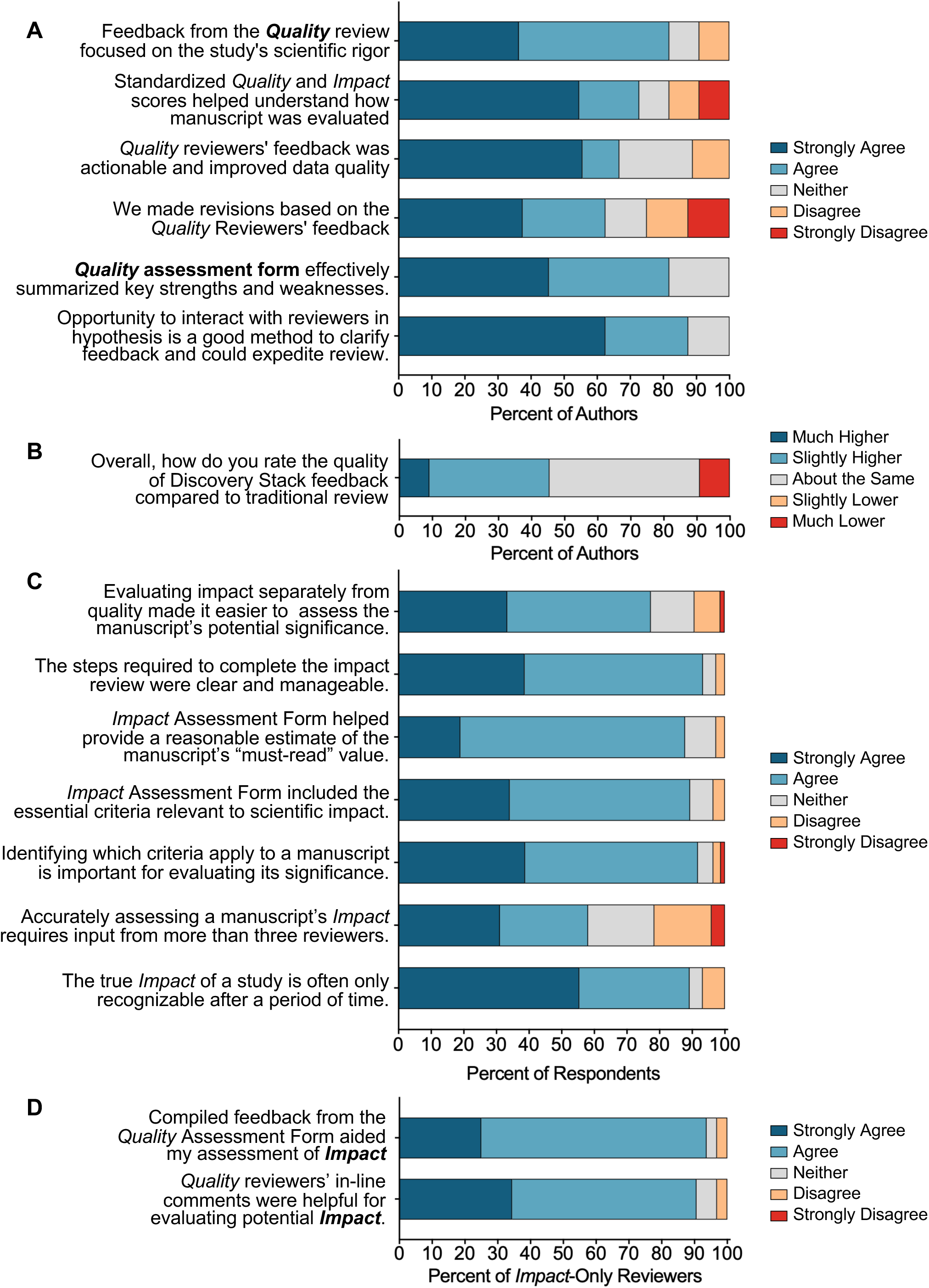
Discovery Stack Enables Rigorous *Quality* Review and Supports Meaningful *Impact* Assessment. (**A**) Authors (n=8-11) were asked to rate their level of agreement with statements related to the feedback they received during the *Quality* review. (**B**) Authors (n=11) were asked to rate the overall usefulness of feedback they received from the Discovery Stack review compared to traditional peer review. (**C**) *Quality/Impact* (n=42) and *Impact*-only (n=32) reviewers responded to questions focused on their experience with the *Impact* assessment. (**D**) *Impact*-only reviewers (n=32) were asked whether the *Quality* reviewers feedback from the assessment form and in-line comments aided their assessment of manuscript *Impact*. The data underlying these figures are available at https://doi.org/10.5281/zenodo.2004122.

When asked to compare the overall Discovery Stack feedback to traditional review, of 11 authors, five rated it better, five rated it similar, and only one rated it worse. (**Fig 8B**). These findings indicate that Discovery Stack delivered rigorous, constructive, and actionable *Quality* reviews. Although limited by a small author sample (n=11), the results support the models’ potential to improve consistency and utility of peer review.

### *Impact* Assessment Viewed as Useful and Strengthened by Prior *Quality* Review

Next, we examined participants’ impressions of the *Impact* review and its value in assessing the broader significance and potential influence of a manuscript’s findings. Most reviewers (77%) reported that evaluating *Impact* independently made it easier to assess significance (**Fig 8C**), consistent with the model’s premise that separating *Quality* and *Impact* enables a more nuanced evaluation.

Reviewers also found the *Impact* review to be well-designed and intuitive; 93% agreed the steps were clear and manageable, 88% found the *Impact* Assessment Form helpful for estimating a manuscript’s “must-read” value, and 89% agreed the form included essential criteria for evaluating significance (**Fig 8C**). Nearly all respondents (92%) agreed that identifying the criteria applicable to each manuscript was essential for an accurate evaluation (**Fig 8C**). Simultaneously, reviewers acknowledged challenges of reliably assessing *Impact*, with 58% agreeing that assessing *Impact* requires more than three reviewers, and 89% agreeing that a study’s true *Impact* often becomes clear only over time, highlighting the importance of dynamic, evolving *Impact* metrics.

To facilitate the separate evaluation, the pilot was designed so that *Quality* reviews were completed before the *Impact* assessment. This allowed *Impact* reviewers to consider *Quality* review feedback while evaluating significance. More than 90% of *Impact*-only reviewers agreed that access to the *Quality* reviews improved their ability to evaluate *Impact* (**Fig 8D**). These findings indicate that reviewers viewed the *Impact* assessment as well-structured, valuable, and strengthened by prior *Quality* review.

### Reviewer Engagement Relies on Outreach, Familiarity, and Trainee Involvement

Recruiting reviewers is a persistent challenge in peer review and a critical barrier to timely evaluations (15, 22). This challenge was amplified for the Discovery Stack Pilot, as most researchers were unfamiliar with the review model and each manuscript required six reviewers. Acceptance rates varied dramatically depending on reviewers’ familiarity with the study and whether they received personal outreach. Among individuals with no prior connection to the pilot, the acceptance rate was only 8.6% (**Fig S5A**). In contrast, reviewers already enrolled in the pilot accepted invitations at a much higher rate of 59% (p < 0.0001, Fisher’s Exact test).

To improve reviewer participation among new reviewers, members of the Scientific Advisory Board (SAB) sent follow-up emails to individuals in their respective networks, which substantially raised the acceptance rate among new reviewers to 53% (**Fig S5A**). Personal outreach increased acceptance odds by more than 12-fold (Odds Ratio (OR)=12.2, *p*<0.0001), while prior enrollment increased odds by 15-fold (OR=15.2, *p*<0.0001). With these strategies in place, for the 17 bioRxiv manuscripts reviewed in the pilot, 283 review invitations were sent (160 for *Quality*/*Impact* and 123 for *Impact*-only reviews), yielding an overall acceptance rate of 33% (**Fig S5B-C**), which is similar to the 32% acceptance rate reported in 2022 by Clarivate’s ScholarOne platform, which supports over 8,000 journals (22). Overall, 90% of accepting reviewers were either familiar with the study or personally contacted by someone in their network. Survey data reinforced this trend, with 70% of reviewers and 64% (7 of 11) of authors reporting they heard about the pilot study prior to participating or alternatively received outreach from a colleague after the initial reviewer request (**Fig S5D**). These findings underscore that outreach and professional networks substantially improve reviewer engagement.

Although trainees who reviewed independently of their advisors represented a small proportion of reviewers (28%; **Fig S5E**), their acceptance rate was markedly higher than that of principal investigators. After adjusting for familiarity and outreach, trainees remained 15 times more likely to accept review invitations (OR=15.3, *p*<0.0001; **Fig S5F**). These findings highlight that actively training and recruiting trainees may be an effective strategy for increasing reviewer engagement.

### Workflow Analyses Identify Operational Barriers to Expedited Peer Review

Another persistent challenge in peer review is the prolonged duration between manuscript submission and publication, which slows the dissemination of scientific findings. Accordingly, an important objective of the Discovery Stack Pilot was to evaluate whether the review workflow could accelerate peer review while maintaining rigorous evaluation. Because the Discovery Stack model included separate *Quality* and *Impact* review phases, total review duration was not directly comparable to traditional journal review. However, the *Quality* review phase was designed to approximate the initial stage of journal review and therefore provided a basis for operational comparison.

The importance of developing more efficient review workflows was underscored by the publication timelines of manuscripts included in the pilot. At the time of the final analysis, 14 of 18 manuscripts had been published in journals. The mean time from submission to publication for manuscripts published in journals was 13 months. The remaining manuscripts were still under review or undergoing revisions, averaging 22 months since submission (**Fig 9A**).

**Fig 9.**
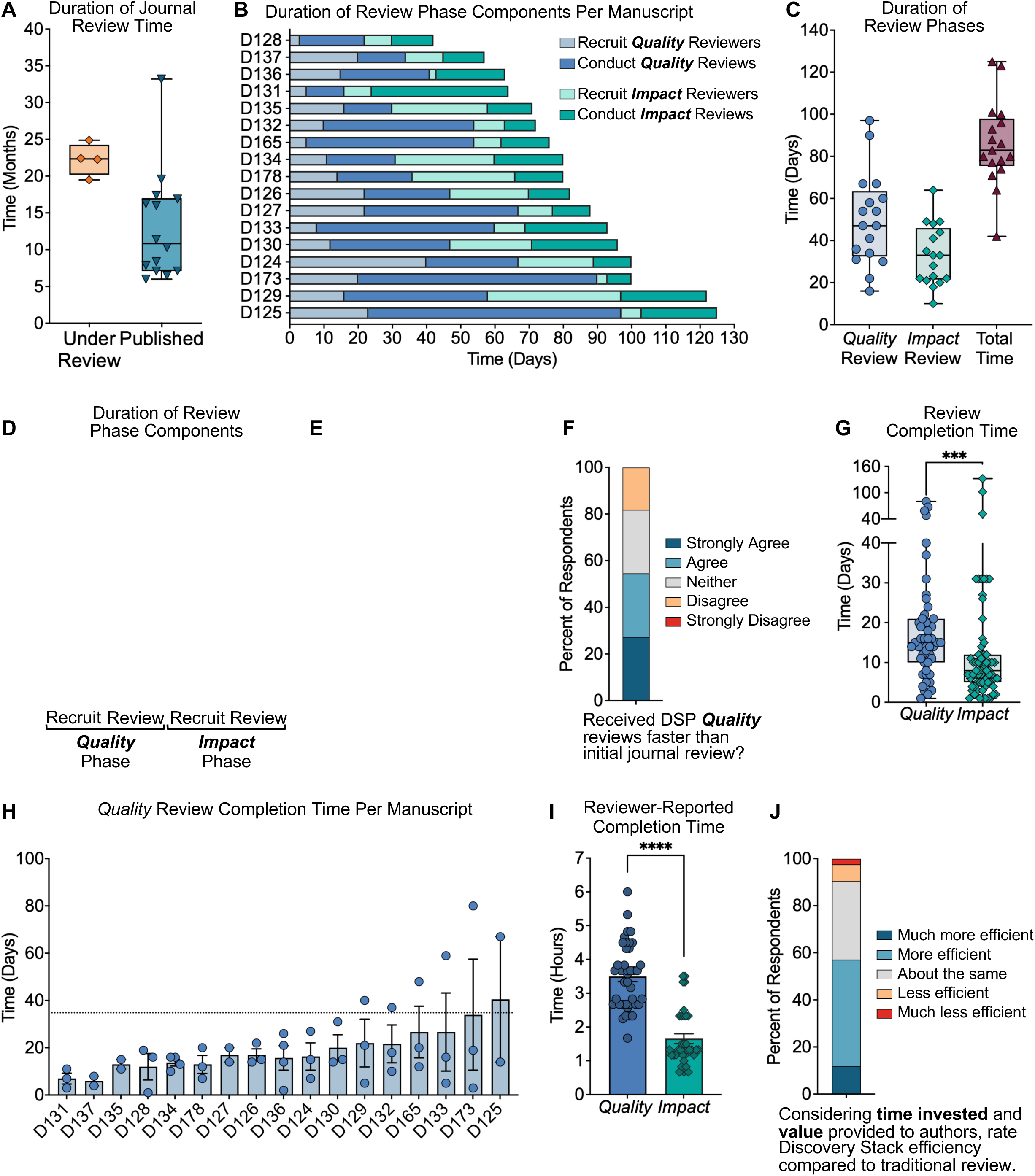
Workflow Analyses Identify Operational Barriers to Expedited Peer Review. (**A**) For manuscripts still under review at the time of the final analysis (08/12/26), the elapsed time from author-reported submission date to analysis date is plotted. For manuscripts published in journals, time from submission to publication reported by the journal is shown. The submission date for manuscript D165 is unknown, so it is excluded from this analysis. (**B**) Days required to recruit three *Quality* reviewers, complete *Quality* reviews, recruit three additional *Impact*-only reviewers, and complete *Impact* reviews is shown for each manuscript. D118 was excluded from analyses in B-D as additional *Impact* reviewers were not recruited due to Hypothes.is compatibility issues. (**C**) Total duration of the *Quality* phase (including recruitment and review completion), the *Impact* phase, and combined review time for each manuscript. (**D**) Average recruitment and review times per manuscript were analyzed with Kruskal-Wallis and Dunn’s multiple comparison post-test. *p*<0.05*; *p*<0.01**. (**E**) Authors (n=11) reported the duration of initial journal review, which was compared to the *Quality* review duration using Wilcoxon matched-pairs signed-rank test. (**F**) Authors (n=11) reported whether they received Discovery Stack *Quality* reviews faster than the initial traditional review. (**G**) Individual *Quality* or *Impact* review times were compared using the Mann-Whitney test (*p*=0.0002***). (**H**) Individual *Quality* Review times were plotted by manuscript. D118 was excluded. (**I**) *Quality/Impact* reviewers (n=40) estimated the time to complete the review, including reading the pre-print, commenting in Hypothes.is, completing both the *Quality* and *Impact* Assessment Forms. *Impact*-only reviewers (n=33) estimated the time to read the pre-print along with *Quality* reviewers’ feedback and complete the *Impact* Assessment Form. (**J**) *Quality/Impact* reviewers (n=42) rated the efficiency of Discovery Stack review, considering both the time spent and value it provided to authors, compared to traditional peer review. The data underlying these figures are available in S1_Table and https://doi.org/10.5281/zenodo.2004122.

Across the pilot, the average time from reviewer recruitment to completion of *both* the *Quality* and *Impact* reviews was 86 days (range: 42–125) (**Fig 9B-C**). The *Quality* phase averaged 50 days and the *Impact* phase 33 days, with reviewer recruitment averaging 15-16 days per phase. The most time-consuming component was completion of *Quality* reviews, which averaged 35 days (**Fig 9D**). Among the 11 manuscripts for which authors reported initial review times, Discovery Stack *Quality* review duration was comparable to the author-reported duration of initial journal review (52 vs. 63 days) (**Fig 9E**). Although the mean Discovery Stack Quality review time was modestly shorter than the author-reported duration of initial journal review, variability between manuscripts was substantial, with several manuscripts receiving Discovery Stack reviews faster than journal review, whereas others were slower. Consistent with this variability, 55% (6 of 11) of authors that completed the surveys reported receiving Discovery Stack *Quality* reviews faster than journal reviews (**Fig 9F**).

To understand why the overall review process was not faster, given that reviewers agreed to complete reviews within 14 days, we examined factors contributing to delays in review completion. Individual turnaround times were generally close to the expected timelines, with *Quality* reviews averaging 18 days and *Impact* reviews 14 days (**Fig 9G**). Moreover, 61% (roughly two out of three) of *Quality* reviewers and 78% of *Impact* reviewers submitted their reviews within three days of the deadline. Visualization of individual reviewer times per manuscript demonstrated that delays were typically due to a single late reviewer rather than widespread delays across all reviewers (**Fig 9H**).

To investigate whether the time required to complete each review contributed to delays in review completion, reviewers were asked to estimate their time investment. *Quality/Impact* reviewers spent an average of 3.5 ± 0.97 hours per review, and *Impact*-only reviewers spent 1.7 ± 0.84 hours (**Fig 9I**). When asked to evaluate the efficiency of the review process, considering both the time invested and the value provided to authors, 91% of *Quality/Impact* reviewers rated it as equal to or more efficient than traditional review (**Fig 9J**).

Together, these findings indicate that streamlining the reviewer workflow alone is insufficient to substantially shorten review duration. Most reviewers adhered to the expected timeline, but isolated late reviews drove the overall delays observed at the manuscript level. Meaningful reductions in review time will therefore require operational interventions specifically targeting the late-reviewer problem, including pre-recruitment of additional reviewers, modest compensation tied to timely submission, and structured contingency plans for replacement when delays exceed defined thresholds.

### Participant Perspectives on Future Publishing Models

To gauge how participants viewed alternative publishing approaches, we surveyed attitudes about needed reforms and desirable platform features. Open-ended responses most frequently identified inefficiencies in the peer review process, unconstructive or biased reviews, and excessive reviewer demands (**Fig S6A-C)**. These concerns aligned closely with the goals of the Discovery Stack model, particularly the use of structured evaluation criteria, separation of *Quality* from *Impact*, in-line commenting, and a peer-improvement mindset.

Participants expressed strong interest in using a new platform that applies scientist-developed metrics to evaluate and curate peer-reviewed research (**Fig 10A**). Respondents also indicated willingness to submit an original research paper to such a platform, but this willingness was lower than using a platform to curate and read research, which likely reflects the continued influence of traditional journals as the key metric of researcher productivity.

**Fig 10.**
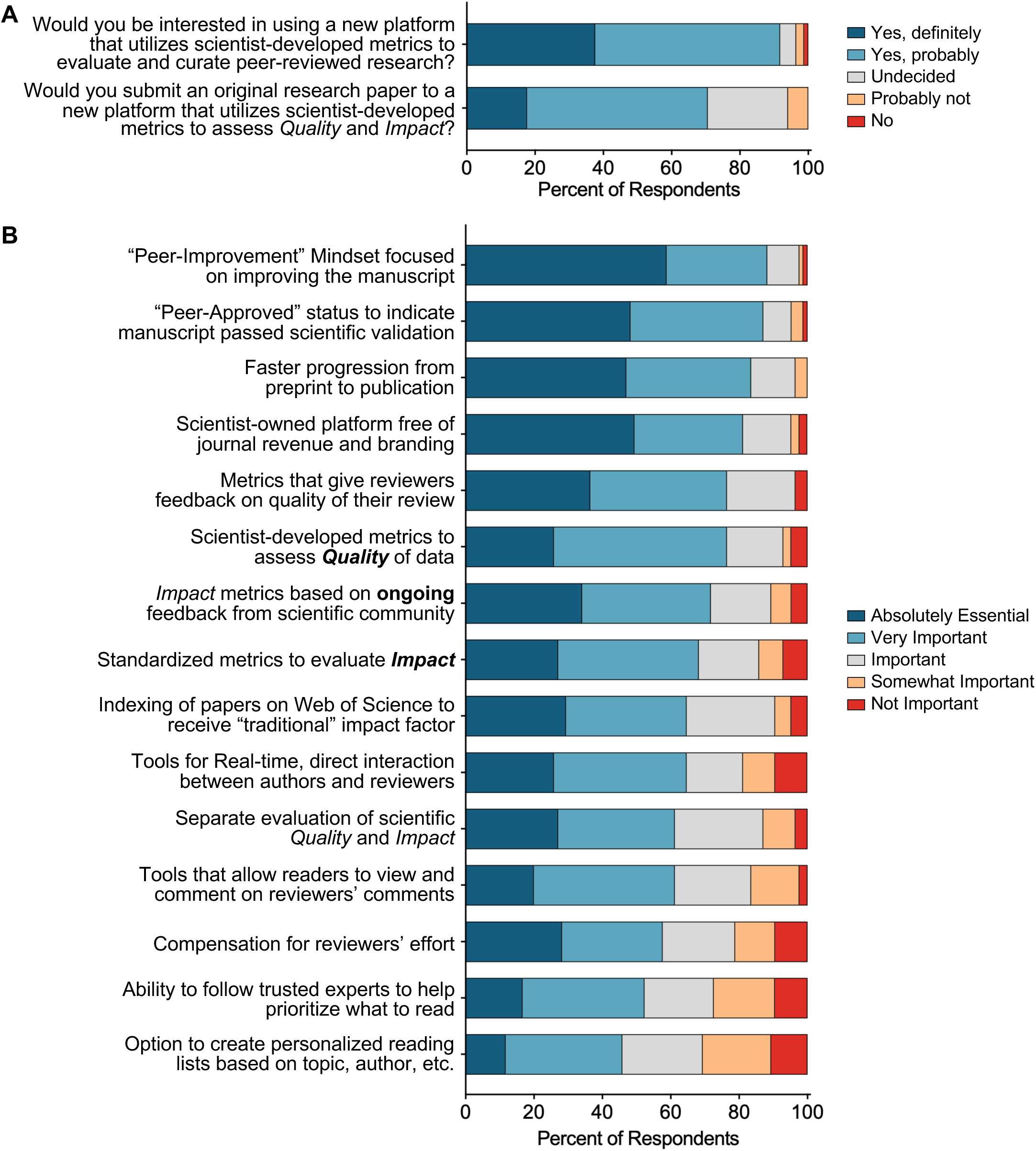
Strong Interest in a New Scientist-Driven Publishing Platform. (**A**) Participants (n=85) were asked whether they would be interested in using a new platform that utilizes scientist-developed metrics to evaluate and curate peer-reviewed research, and whether they would submit an original research paper to a new platform that utilizes scientist-developed metrics to assess *Quality* and *Impact*. (**B**) Participants (n=85) were asked to rank the importance of features that would influence their likelihood of using a novel platform for evaluating and curating peer-reviewed and peer-approved research. Features were rated from “absolutely essential” to “not important”. The percent of respondents selecting each rank is shown. The data underlying these figures are available at https://doi.org/10.5281/zenodo.2004122.

When asked which features would most influence adoption of a new platform, participants prioritized a “peer-improvement” mindset (88%), a “peer-approved” status for validated manuscripts (87%), faster publication timelines (84%), a scientist-owned platform free of journal revenue and branding (81%), reviewer feedback metrics (77%), scientist-developed *Quality* metrics (77%), and evolving *Impact* metrics based on *ongoing* community input (72%) (**Fig 10B**). Additional requested features included institutional recognition by funding and promotion committees, improved access to underlying data and code, and moderation of comments to maintain constructive review *Quality*.

Given the modest cohort size and potential for selection bias towards participants receptive to alternative peer review models, these findings should be interpreted cautiously. Nevertheless, they suggest that several of the core features tested in the Discovery Stack Pilot align with researchers’ priorities and highlight the need for institutional recognition of the platform to overcome hesitancy to depart from traditional journals.

## DISCUSSION

The Discovery Stack Pilot evaluated the feasibility and value of a scientist-designed peer review model that separates scientific rigor (*Quality*) from perceived significance (*Impact*), uses standardized metrics, incorporates in-line commenting, and promotes a peer-improvement mindset. Conducting reviews in parallel with traditional journal reviews enabled a direct comparison of feasibility and effectiveness. Several important findings emerged. Notably, reviewers successfully evaluated *Quality* distinctly from *Impact*, supporting the conceptual separation of these dimensions. *Impact* assessments showed greater variability than *Quality* assessments, highlighting the importance of broader reviewer input when evaluating scientific significance and reinforcing the rationale for independently evaluating these dimensions. Participants also strongly endorsed core elements of the model, including structured evaluation criteria, standardized metrics, and the separation of *Quality* from *Impact*. Together, these findings support the feasibility of the Discovery Stack framework and provide a foundation for larger-scale evaluation.

A major outcome of this pilot was empirical support for separating *Quality* from *Impact*, as we had previously proposed separating these two dimensions to address the long-standing problem that perceptions of significance can mask concerns about rigor, and vice versa (21). Although *Quality* and *Impact* were positively associated, as expected, when poor rigor limited perceived importance, residual and tertile analyses revealed frequent divergence. Moreover, *Impact* scores, but not *Quality* scores, significantly correlated with JIFs. Although this observation should be interpreted cautiously given the pilot study’s modest sample size, it is consistent with growing evidence that JIF is a poor proxy for the methodological rigor and reliability of individual studies (11). JIFs also have important conceptual and practical limitations as measures of the quality or impact of individual publications (9–14). Because citation distributions are highly skewed, the arithmetic mean used to calculate JIFs may not accurately represent the citation performance of a typical article within a journal (12–14). Additionally, publishers can negotiate which article types are included in the denominator, with independently verified discrepancies of up to 19% compared to Clarivate’s published figures (10). Importantly, studies suggest that journals with higher JIFs do not consistently publish more reliable research (10, 14). For example, higher JIFs are associated with higher retraction rates, although this relationship may partly reflect the greater visibility and scrutiny of articles published in highly cited journals (10, 14). Data from the Discovery Stack Pilot demonstrate that reviewer-assessed *Quality* captured information distinct from conventional journal-based JIFs, which supports the rationale for evaluating scientific rigor independently of perceived significance. Importantly, most reviewers (93%) agreed that the Discovery Stack model effectively supported separate assessments, and 85% found that separating these dimensions led to more constructive and insightful reviews. Separating *Quality* and *Impact* would substantially benefit science by assigning value to high-*Quality* work, independent of novelty, creating space for careful replication studies, incremental clarifications, and contrarian findings that the current novelty-oriented model often overlooks.

Data from the pilot also revealed a greater dispersion in *Impact* scores relative to *Quality*, reflecting inherent subjectivity in evaluating significance, which is sensitive to field coverage, methodological preferences, translational focus, and individual research agendas. Operationally, this finding supports the use of a larger number of *Impact* reviewers per manuscript to capture a representative distribution of viewpoints.

The pilot also evaluated the feasibility and utility of standardized metrics. Participants strongly endorsed metrics for both *Quality* (90%) and *Impact* (82%). This support likely reflects not only the resulting quantitative scores but also endorsement of the structured framework itself, including the use of clearly defined evaluation criteria, which is uncommon in traditional peer review. In addition to providing data to generate metrics, the Likert-scale questions aligned reviewer attention to the core elements of rigor (methodology, controls, statistics, concordance between data and conclusions) or significance (extent of advance in scientific understanding, filling of a knowledge gap, “must-read” value), supporting more consistent evaluation across manuscripts.

One potential long-term application of standardized *Quality* metrics would be the development of empirically derived thresholds for scientific rigor. With substantially larger datasets and appropriate validation of score calibration, reproducibility, and predictive performance, it may become possible to identify score ranges that reliably distinguish manuscripts meeting predefined standards of scientific rigor. Although the present pilot was not designed to establish such thresholds, it demonstrates the feasibility of generating standardized *Quality* scores that could serve as the foundation for future investigation. Like traditional peer review, revised manuscripts would undergo additional rounds of evaluation, with updated *Quality* and *Impact* scores updated following each revision. The final *Quality* score could serve as an indicator of rigor and reliability of the evidence supporting a manuscript’s conclusions.

In contrast, because scientific significance evolves as discoveries are replicated, extended, and applied, an important long-term goal of the Discovery Stack framework would be the development of dynamic *Impact* metrics rather than fixed assessments at the time of publication. Such metrics could integrate three complementary sources of information: 1) expert assessments of significance generated at the time of review, 2) citation data from bibliometric databases, capturing formal scientific uptake, and 3) broader indicators of influence, including readership, downloads, reference manager saves, press coverage, policy citations, and social media engagement. This framework would require rigorous empirical evaluation of its strengths and limitations as each of these components have limitations. While overreliance on metrics alone risks oversimplification (23), these hazards can be mitigated when metrics are based on clearly defined attributes, coupled with narrative feedback and in-line annotations, and routinely evaluated.

The Discovery Stack model builds upon several recent innovations in peer review that seek to improve the evaluation and communication of scientific research (24). Soundness-only peer review, pioneered by journals such as PLOS ONE and later adopted by other journals, aims to evaluate methodological rigor while leaving judgments of novelty and significance to the broader scientific community after publication. However, studies suggest that reviewers and editors often continue to incorporate traditional assessments of importance and novelty into their recommendations, highlighting the practical difficulty of separating these dimensions in a conventional editorial framework (24). The Discovery Stack model addresses this challenge by explicitly evaluating and scoring both dimensions using defined criteria.

The publish-review-curate model, implemented by eLife and now used on other platforms (25), represents another important innovation by distinguishing the strength of evidence from the significance of findings, while emphasizing transparent evaluation after publication. The Discovery Stack model shares this objective but extends the concept by generating standardized *Quality* and *Impact* metrics that allow readers to filter and prioritize work by rigor and significance at scale, rather than evaluating manuscripts on a journal-by-journal basis.

Open review models represent another innovation now used by many journals to increase transparency by publishing reviewer comments alongside manuscripts. While this approach provides valuable context and insight into the review process, readers must still evaluate individual reviews to identify rigorous and influential studies. As the volume of published research continues to grow, this becomes increasingly challenging. In addition, some journals have incorporated in-line annotation tools during peer review of manuscripts, demonstrating the feasibility of integrating contextual comments directly within manuscripts (26). The Discovery Stack model builds upon existing approaches by combining transparent review, independent *Quality* and *Impact* assessments, in-line annotation, metrics based on standardized criteria, and a framework for dynamic post-publication evaluation of scientific influence. Together, these features enable readers to filter, search, and prioritize manuscripts by rigor and significance at scale, rather than relying on journal prestige or reading the full text of every review.

Registered Reports are another innovation that seeks to improve research rigor by evaluating study design before results are known, reducing publication and reporting biases (27). This approach is particularly well suited to hypothesis-driven studies with predefined experimental protocols, whereas the Discovery Stack model provides a complementary framework for evaluating the *Quality* and *Impact* of completed research across a broad range of study types.

Timeliness remains one of the most significant challenges in peer review. Despite emphasizing deadlines and streamlining reviews by using in-line comments instead of lengthy narratives, the Discovery Stack *Quality* review phase averaged about eight weeks, matching the initial journal review rather than shortening it as intended. Individual reviewers generally adhered to the expected timelines, and most delays were caused by a single reviewer. Replacing a late reviewer rarely shortens the overall review time because confirming a replacement takes time, and the new reviewer requires time to complete the review. While many journals address this by relying on two reviewers instead of three, this approach risks weakening the overall *Quality* of the evaluation. The Discovery Stack Pilot demonstrated that reducing the review duration requires more than workflow simplification. Rather, clear accountability and pre-planned backup coverage are required. Practical steps include confirming four reviewers at the outset, recruiting “alternate” reviewers, and rewarding timely submissions with monetary compensation and recognition. Notably, a recent study combining pre-recruitment of expert reviewers, compensation, and strict deadlines achieved review turnaround times of less than one week, illustrating how innovative recruitment strategies and incentives can expedite the review process (28).

Reviewer recruitment and reviewer burden are practical considerations for adoption of the Discovery Stack model, as reviewer recruitment remains a major bottleneck in peer review and adds substantially to total review time (15, 22). In our pilot, confirming three *Quality* reviewers averaged two weeks per manuscript, a nontrivial delay that accounted for one quarter of the eight-week *Quality* review phase. Rather than recruiting nine fully independent reviewers per manuscript (three *Quality* and six *Impact*), *Quality* reviewers also performed a separate *Impact* assessment of the same manuscript. Subsequently, three additional *Impact*-only reviewers were recruited specifically to increase the number and diversity of *Impact* evaluations. Importantly, the *Impact*-only review phase was designed to reduce reviewer burden by focusing exclusively on scientific significance and allowing reviewers to reference prior *Quality* reviewer comments addressing rigor and methodology. Consistent with this streamlined role, *Impact*-only reviewers reported spending an average of 1.7 hours per review, substantially less than the 3.5 hours reported by reviewers to complete the *Quality* assessment. Thus, additional *Impact* reviewers add less to the total reviewer burden than their number alone would suggest. Nevertheless, reviewer recruitment remains an important consideration for future implementations of the model. Reducing the reviewer bottleneck will ultimately depend on increasing the reviewer pool. Our recruitment analyses provide important guidance for future deployment of the Discovery Stack model. Familiarity with the project and direct outreach markedly improved reviewer acceptance rates, suggesting that current enthusiasm for the model may not yet extend broadly across the wider research community and that broader implementation will require strategies to engage scientists beyond early adopters and existing professional networks. Additionally, trainees were especially responsive to accepting review assignments. These insights suggest that visible recognition, modest compensation, and formal training for early-career scientists may help expand the reviewer pool, while also providing opportunities to develop constructive peer review skills and establish scientific reputations. Modest reviewer compensation was intentionally incorporated into the pilot because equitable compensation for peer-review labor is a central principle of the Discovery Stack model. However, the present study was not designed to determine the relative contributions of compensation, professional outreach, prior familiarity, and scientific interest to reviewer participation. Notably, 75% of compensated reviewers (56 of 75) elected to donate their honoraria rather than accept it, suggesting that financial compensation was not sole motivation for many participants. Nonetheless, identifying sustainable funding sources to support for reviewer compensation will be an important consideration for implementation of alternative peer-review models.

Despite recruitment challenges, survey responses from newly recruited participants were broadly similar to those from previously engaged participants, suggesting that support for the Discovery Stack model was not solely driven by prior familiarity with the project. Nevertheless, because participation was voluntary and recruitment relied on existing networks, these findings should be interpreted cautiously. Within this cohort, revealed broad dissatisfaction with current publishing norms and strong enthusiasm for reform. Accordingly, 92% of participants indicated a willingness to use a scientist-designed platform to evaluate and curate peer-reviewed research. Features most likely to drive adoption align with the Discovery Stack model, including a peer-improvement mindset, a “peer-approved” status to indicate scientific validation, faster timelines to publication, scientist ownership, and reviewer feedback metrics. These results suggest the Discovery Stack model provides a promising foundation for further refinement and evaluation.

Although willingness to submit manuscripts was strong (71%), it lagged behind readiness to use a platform for evaluation and curation, reflecting continued dependence on journal prestige and JIFs for career advancement and funding decisions. Despite the limitations of JIFs, these metrics remain the primary mechanism used by funding and academic institutions to assess the value and impact of scientific research. The reliance on journal-based JIFs presents a major barrier to the adoption of new publishing models. Accordingly, successful implementation of scientist-designed peer review platforms will require not only technical innovation but also broader acceptance by funding agencies, academic institutions, and other stakeholders. Encouragingly, movements promoting improved research assessment are emerging (7, 23, 29), and the Discovery Stack model, with its independent evaluation of scientific rigor and impact, may provide a practical complement to these initiatives.

This study has limitations. The modest sample size limits statistical power and the strength of survey-based conclusions. The focus on immunology and cancer biology preprints, selected from a predefined eligibility pool rather than randomly sampled across bioRxiv, limits generalizability of the findings to other scientific disciplines. In addition, the pilot was not designed to incorporate explicit stratification by manuscript or author characteristics, including geographic distribution, institutional diversity, or other demographic features. Because participation was voluntary and leveraged prior engagement and professional networks, the study population may be enriched for participants more receptive to alternative peer review models, which further constrains generalizability of participant perceptions. In addition, surveys were completed by a single representative author from each participating manuscript, typically the senior author or first author responsible for the review process. Consequently, this pilot was not designed to evaluate whether perceptions differed according to authorship role or career stage. Future studies should include larger numbers of authors from individual research teams to determine whether perspectives vary among senior authors, first authors, and other contributors. Other limitations include incomplete publication outcomes for some manuscripts, reliance on an external annotation tool, the evaluation of mostly initial submissions, which limited both the analysis of score dynamics across revisions and direct comparison of Discovery Stack scores with the JIFs of journals in which manuscripts were ultimately published. These constraints are typical of feasibility studies and highlight the need to address these limitations in future iterations through larger and more diverse cohorts.

In summary, the Discovery Stack Pilot demonstrated that reviewers reliably evaluated *Quality* and *Impact* as separate dimensions, and that *Impact*, by its nature, varies more than *Quality*. The model’s core elements of standardized metrics, in-line annotation, peer-improvement mindset, and distinct *Quality* and *Impact* evaluations enhanced clarity and constructiveness, earning considerable support. Overall, these results provide a practical foundation for continued refinement and future studies to assess the performance, scalability, and broader applicability of the Discovery Stack model across more diverse scientific communities.

## MATERIALS AND METHODS

### Study Design

The Discovery Stack Pilot consisted of three sequential phases designed to test a structured, peer review process that separately evaluated *Quality* and *Impact*. These phases were: 1) *Quality* Review, 2) Author Response, and 3) *Impact* Review.

### *Quality* Review Phase

The *Quality* review phase provided a structured, standardized evaluation of each manuscript’s scientific rigor. Reviewers were instructed that *Quality* referred to the appropriateness of experimental design, methodology, controls, statistical analyses, and sample sizes, as well as whether the data supported the stated conclusions. To support consistency and emphasize that improving scientific *Quality* was the primary objective of this phase, reviewers received a *Quality Assessment Checklist* (**Fig S7**) directing them to evaluate four key areas: 1) whether conclusions were adequately supported by the data and free of contradictions, 2) soundness of the experimental design, controls, methodology, and statistical analyses, 3) data integrity and reproducibility, including sample sizes and number of replicates, and 4) clarity of presentation.

*Quality* reviewers provided feedback through two complementary mechanisms. First, in-line comments were added directly to the manuscript preprint using the Hypothes.is tool, mirroring how scientists typically provide feedback to colleagues during manuscript preparation. Comments were initially posted in private groups visible only to the editor, then compiled and shared in groups visible to all reviewers and the authors. Comments from reviewers who wished to remain anonymous were de-identified before sharing.

Second, reviewers completed a standardized *Quality* Assessment Form consisting of six short-answer questions and multiple Likert-scale questions (**Fig 2A-B**). This form was designed to capture summary-level assessments and test the feasibility and utility of using structured, quantitative metrics to evaluate scientific rigor. All assessment forms were designed, distributed, and collected using the Formaloo platform.

### Author Response Phase

Following submission of *Quality* reviews, feedback was compiled and shared with authors and other reviewers. In-line comments were shared by inviting authors and reviewers to a shared Hypothes.is group containing all reviewer annotations. *Quality* Assessment Form feedback was compiled in a summary PDF that included short-answer responses and graphical representations of the Likert-style questions (**Fig 2C**). Both in-line comments and assessment form feedback were labeled by reviewer. Reviewers who disclosed their identity were named, while anonymous reviewers were designated “Reviewer #”. Authors were encouraged, though not required, to reply directly to reviewers in Hypothes.is. Authors were not required to conduct additional experiments for the pilot.

### *Impact* Review Phase

*Impact* was defined as the extent to which a study advances scientific understanding, addresses critical knowledge gaps, influences multiple fields, or has therapeutic relevance. Reviewers were informed that a manuscript could be of high *Quality* yet have limited *Impact* if its contribution was incremental or relevant to only a small audience.

To ensure sufficient evaluations for each manuscript, *Quality* reviewers were also asked to provide a separate assessment of the manuscript’s potential *Impact*. After completing the *Quality* Assessment Form, reviewers were directed to a separate *Impact* Assessment Form including questions evaluating transformative potential, generalizability, technological advancement, and mechanistic insight (**Fig 3A-D**). This structured framework enabled *Impact* to be evaluated independently of *Quality* while capturing the diversity of perspectives on scientific significance.

### *Impact*-only Review Phase

Since assessment of *Impact* is inherently subjective, additional reviewers were recruited to focus solely on the *Impact* evaluation. The goal was to obtain a minimum of three *Quality* and six *Impact* reviews per manuscript. *Impact*-only reviewers were recruited after the *Quality* phase was completed and they received the manuscript, *Quality* reviewers in-line comments, and assessment form feedback. *Impact*-only reviewers completed the same *Impact* Assessment Form described above (**Fig 3A-D**).

### Reviewer Experience, Training, Expectations, and Compensation

Reviewers were selected based on their subject matter expertise, identified through author recommendations, nominations by members of the scientific advisory board, previously enrolled participants, or PubMed searches by the editor. While most reviewers were principal investigators, trainees also completed reviews with prior endorsement from their PI or as part of collaborative reviews. Most reviewers brought substantial experience to the process: 83% previously reviewed 10 or more manuscripts, 49% reported over a decade of experience submitting and reviewing papers, and 80% had been engaged in peer review for at least five years (**Fig S5G-H**).

Before receiving review materials, reviewers were invited to attend a brief Zoom onboarding session or receive detailed instructions via email. Most *Quality* reviewers (88%) attended a Zoom onboarding meeting, whereas most *Impact*-only reviewers (90%) received detailed instructions via email. Onboarding covered Hypothes.is setup, pilot goals, and emphasized the value of a peer-improvement mindset, defined as offering clear, constructive, and actionable feedback to improve scientific rigor and suggest additional experiments only when essential to support the conclusions, feasible for the research group, and within the scope of the study. *Quality* reviewers agreed to complete reviews within 14 days and *Impact*-only reviewers within 10 days. Timeliness was communicated during reviewer recruitment, reinforced during onboarding sessions, emphasized in the review instructions, and reiterated in follow-up reminder emails. To acknowledge their contributions and reflect our commitment to a platform that compensates reviewers, *Quality* reviewers were offered $30, and *Impact*-only reviewers were offered $20 per review. Reviewers had the option to donate their compensation to Solving for Science rather than accept payment. Of the 75 reviewers who completed the surveys, 55 (73%) elected to donate their compensation.

### Manuscript Enrollment

Five manuscripts were initially enrolled following email invitations to previously enrolled participants. Subsequently, manuscripts were recruited from recent bioRxiv postings based on predefined criteria, including 1) relevance to immunology or cancer biology and 2) recent posting date. Using these criteria, 118 authors were contacted, resulting in the enrollment of 11 more manuscripts. Two additional bioRxiv preprints were included in the pilot despite lack of author response, as reviewers with appropriately matched expertise had already been identified and agreed to participate. In total, 18 manuscripts were reviewed in the Discovery Stack Pilot. One manuscript (D118) was hosted on SSRN rather than bioRxiv and was excluded from some analyses due to incompatibility of Hypothes.is and SSRN.

At the conclusion of the pilot, 11 of 18 authors (61%) completed the author feedback survey, which captured information about the submission status of their manuscripts. Of these 11 manuscripts, eight were first-time submissions to a traditional journal, while three were revised drafts following one round of journal revision. As of August 12, 2026, 14/18 manuscripts reviewed in the pilot had been accepted for publication in a traditional journal. Upon completion of the pilot, surveys were sent to a single author from each manuscript, either the senior author or the first author as middle authors are typically less involved in the review process. Nine of the 11 responding authors were principal investigators and senior authors, while the remaining two author respondents were first authors (a postdoc and graduate student).

### Development and Validation of Composite *Quality* and *Impact* Metrics

*Quality* and *Impact* metrics were developed using standardized assessment forms containing Likert-scale questions rating defined attributes of each dimension on a 1-5 scale (1= strongly agree, best; and 5=strongly disagree, worst (**Fig 2-3; S1A-D**).

*Quality* scores were derived from 13 Likert-scale questions addressing experimental design, controls, statistical analysis, reproducibility, and data support for conclusions. Each question was weighted according to its relative importance, and weighted responses were averaged to yield a single composite *Quality* score per review (**Fig S1A**). All manuscripts received at least two composite *Quality* scores, most received three. To evaluate the effect of weighting, weighted and unweighted averages were compared to the average score of a key item (“Overall, the data support the conclusions presented in the paper”), which served as a benchmark of overall rigor. Weighted scores aligned with unweighted averages but trended toward the benchmark (**Fig S1C**).

*Impact* scores were generated from one multiple-choice question and five Likert-scale items. In the multiple-choice question, reviewers selected features contributing to a manuscript’s *Impact* (**Fig 3**). Each feature was weighted by importance, and the weighted sums were normalized to a 1-5 scale and combined with the weighted average of the five Likert-scale responses to generate a single composite *Impact* Score for each review. All manuscripts received at least five composite *Impact* scores, and most (15 of 17) received six or more. Unweighted and weighted averages, with and without the multiple-choice question, were compared and showed similar results (**Fig S1D**).

Most manuscripts reviewed in the pilot were initial submission. However, three manuscripts were revised versions after one round of review. Composite *Quality* and *Impact* scores of initial and revised submissions were compared to determine if revised manuscripts should be analyzed separately. No significant differences, or even trends toward higher scores were observed in the revised manuscripts, so all manuscripts were grouped together for further analyses (**Fig S1E-F**).

### Composite *Quality* and *Impact* Score Analysis

Associations between *Quality* and *Impact* scores were tested using Pearson’s correlation and linear regression models in GraphPad Prism. Analyses were performed using all available *Quality* and *Impact* scores, and separately using only scores from reviewers who evaluated both *Quality* and *Impact*.

Variability between *Quality* and *Impact* scores was assessed using standard deviation (SD), range, and interquartile range (IQR). Quartiles were calculated using the inclusive quantile definition (QUARTILE.INC in Excel) and IQR was defined as the difference between the 75^th^ and 25^th^ percentiles (Q3-Q1). Differences were compared using the Wilcoxon matched-pairs signed-rank test. Manuscript D136 was excluded as an outlier (z-score = −3.14, >3 SD from mean difference). To account for unequal reviewer numbers, a subsampling approach was implemented using R. For each manuscript, *Impact* scores were randomly subsampled to match the number of *Quality* scores, and variability metrics were recalculated for each subsampled dataset. The subsampling procedure was repeated for 5,000 iterations to generate empirical distributions of the mean differences (*Impact* - *Quality*) for each variability metric. Confidence intervals and one-sided p-values were derived from these distributions, with p-values defined as the proportion of iterations in which the mean difference (*Impact* - *Quality*) was less than or equal to zero.

### Survey Design and Analysis

At the completion of the pilot study, surveys were distributed to 93 individuals who participated as authors, *Quality/Impact* reviewers, or *Impact*-only reviewers. Author surveys were distributed only to the author who enrolled the manuscript in the Discovery Stack Pilot, as these individuals were directly engaged with the Discovery Stack review process and this approach avoided disproportionate weighting of manuscripts with large author lists. Surveys were generated and distributed using the Formaloo platform. Three distinct surveys were developed, each tailored to the participant’s role. Questions on each survey are available at https://doi.org/10.5281/zenodo.20041225. Individuals who served in multiple roles received a separate survey for each role. In total, 101 survey invitations were sent, and 86 completed responses were received, yielding an 85% completion rate.

Responses included a mix of Likert-scale, multiple-choice, and open-ended formats. Quantitative responses were summarized as percentages of total respondents per question. Open-ended responses were coded thematically, and frequencies were tallied to identify common themes.

To assess potential bias associated with prior familiarity with the study, responses from participants new to the study (NEW) were compared to those from previously enrolled participants (DSP-enrolled) using Mann-Whitney U tests. Analyses were restricted to survey items with sufficient responses across groups. Questions that were only administered to authors were excluded due to the limited number of author respondents (NEW, n = 7; DSP-enrolled, n = 4). For all other questions, responses from all participant roles (*Quality/Impact* reviewers, *Impact*-only reviewers, authors) were pooled within the NEW and DSP-enrolled groups and compared. Lower scores corresponded to more favorable responses. P-values were adjusted using the Benjamini–Hochberg procedure. Effect sizes (r) were calculated from the standardized Mann-Whitney statistic as r = Z/√N, where Z was derived using the normal approximation with continuity and tie corrections and N is the total number of observations. These analyses were performed in R.

## Data, Code, and Software Availability

Statistical analyses were performed in GraphPad Prism (version 11.0.1) or R (version 4.5.1 (2025-06-13) with RStudio (version 2026.04.0+526). Surveys and assessment forms were administered using the Formaloo platform (https://www.formaloo.com). Figures were generated in Prism or R and assembled in PowerPoint. Source data are available in Supplementary Tables or at https://doi.org/10.5281/zenodo.20041225, as indicated in the Figure Legends. The code for analyses is available at https://github.com/mmcgargi/Discovery-Stack-Analysis.

## AUTHOR CONTRIBUTIONS

MAM: Conception of methods, coordination of study, data analysis and presentation, writing and editing of manuscript.

BCL: Conception of study, ongoing coordination during the study, initial setup of study.

KC, CR, MK, MK, BR, SS, AG, LBR, SR, TS, IR, JG, JO-M, SS, DM, AO, IB, HGV, JS, TF, NJ: Conception of study, ongoing coordination during the study.

MFK: Conception of study, ongoing coordination during the study, writing and editing of manuscript.

## ACKNOWLEDGEMENTS

We thank Solving For Science for funding and support associated with undertaking this study. We thank the NIH for respective funding for all the labs involved in this study. We thank Igor Brodsky, Julien Gaillard, Ananda Goldrath, Nikhil Joshi, Savan Ram, Carla Rothlin, Sunny Shin, Sara Suliman, and Joe Sun for helpful advice during the planning, implementation, and analysis of the pilot.

## SUPPLEMENTARY INFORMATION

**Fig S1.**
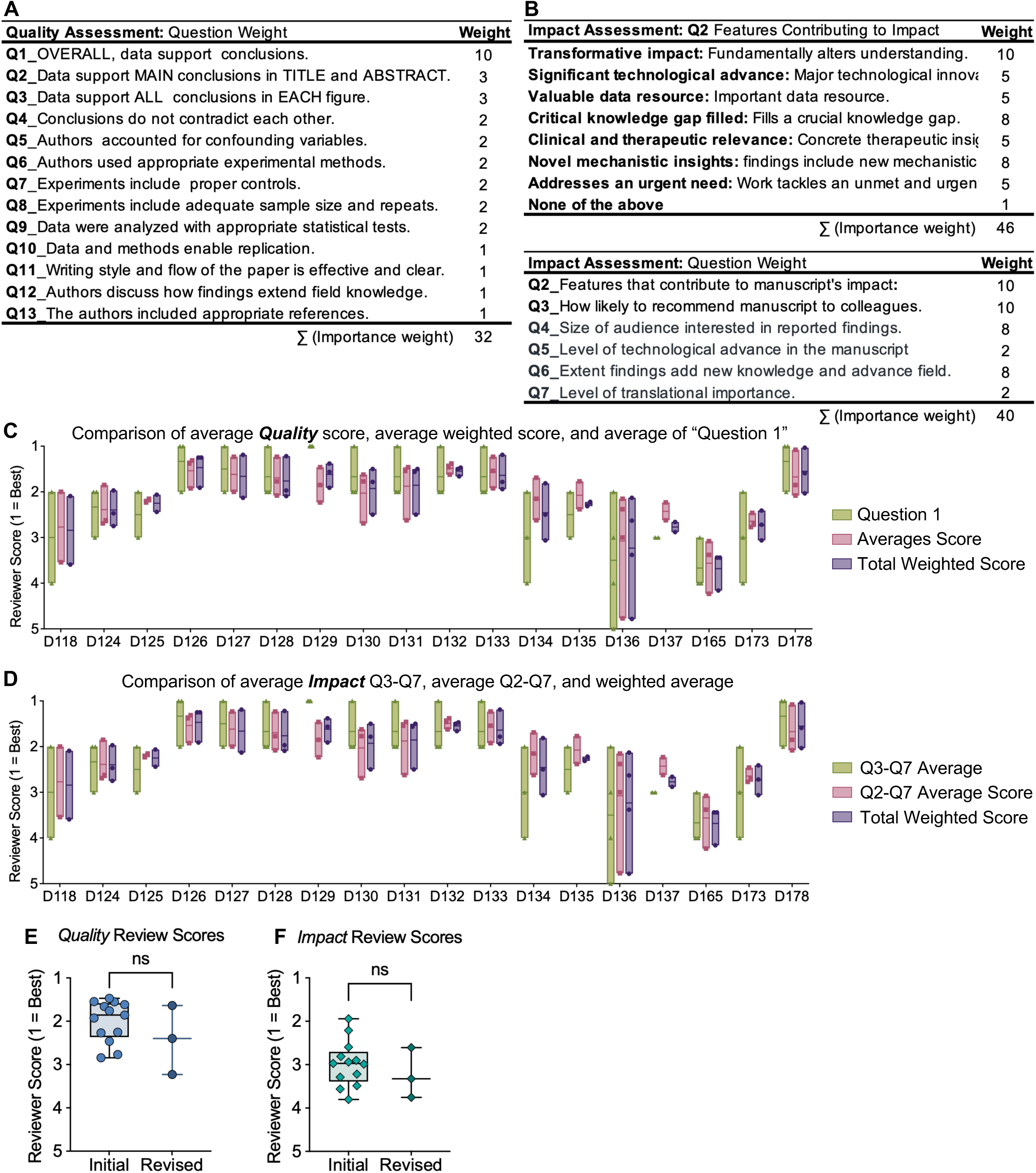
Calculation and Comparison of *Quality* and *Impact* Composite Scores. Composite *Quality* and *Impact* review scores were calculated to condense the full set of ratings from each review into a single metric. (**A**) Composite *Quality* scores were derived from 13 Likert-scale questions that were weighted based on relative importance. Composite *Quality* score = ∑ (Question Score x Importance weight)/ ∑ (Importance weight). (**B**) Composite *Impact* scores were based on responses to one multiple-choice question and five Likert-scale items. To integrate responses across two question types, each feature in the multiple-choice question was assigned a weight reflecting its relative importance to *Impact*, the weighted sum of selected features was calculated and then normalized to a 1-5 scale using a linear transformation. Transformed Q2 = 6 - (1 + ((∑Q2-1)*4/(36–1))). In this scale, selecting no features corresponds to a score of 5 (lowest *Impact*) and selecting features totaling the maximum score achieved (36) corresponds to a score of 1 (highest *Impact*). This transformed Q2 value was then combined with the weighted average of the five Likert-scale responses to generate a single composite *Impact* score for each review. (**C**) The weighted *Quality* composite score, unweighted average of Likert-scale responses, and the average score for the first statement (Overall, the data support the conclusions presented in the paper) (**D**) The weighted composite *Impact* score (including Q2), the unweighted average of Q2-Q7, and the unweighted average of Q3-Q7 were compared across manuscripts. (**E-F**) Weighted composite scores for (**E**) *Quality* or (**F**) *Impact* were compared between manuscripts enrolled as initial submissions versus revised versions. The version of three manuscripts (D165, D173, D178) was unknown, so are excluded. No significant differences were observed by Mann-Whitney test. The data underlying these figures are available at https://doi.org/10.5281/zenodo.2004122.

**Figure S2.**
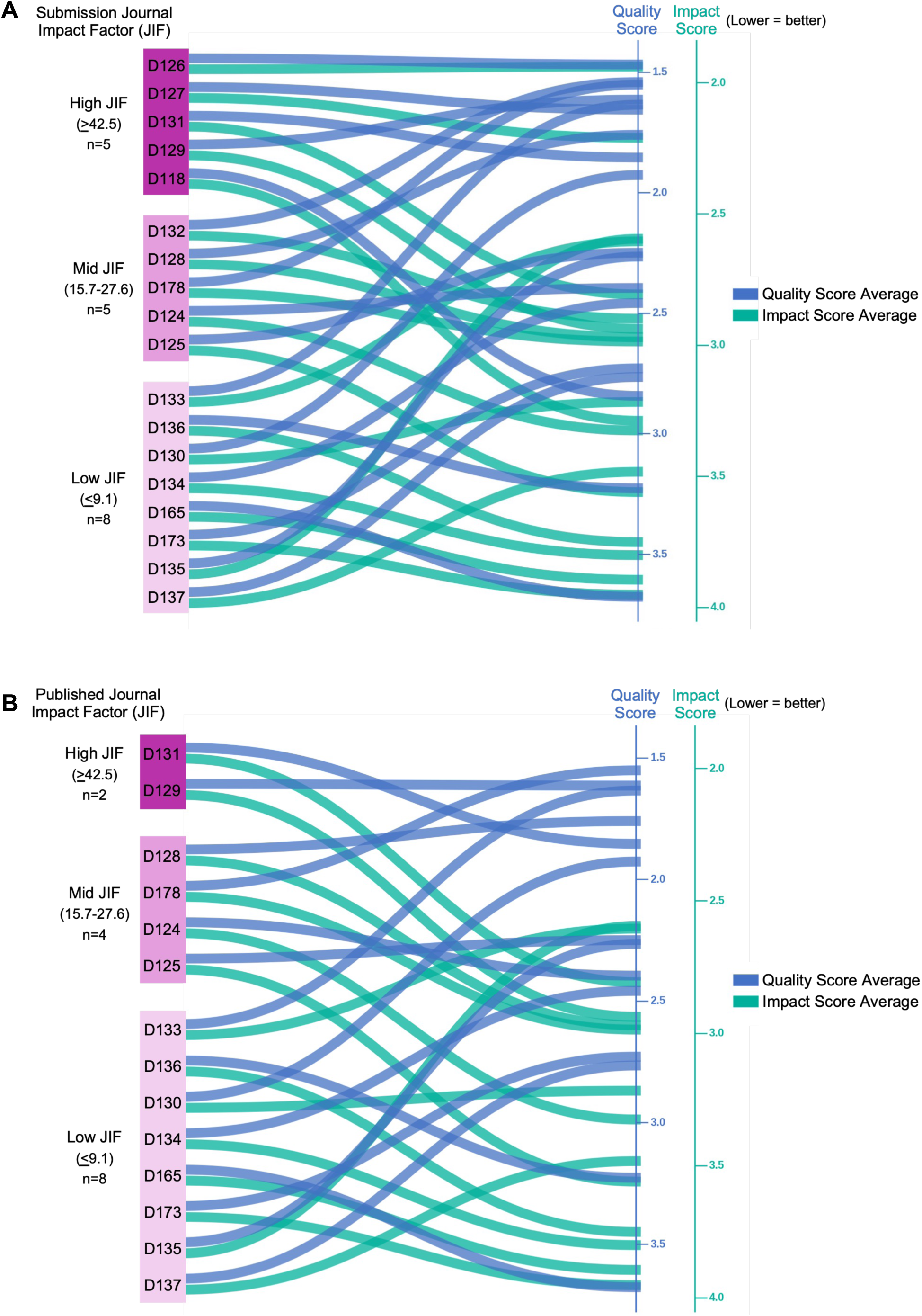
Alluvial plot of reviewer *Quality* and *Impact* scores by submitted JIF tier. Manuscripts are grouped on the left by the JIF of the journal to which it was (**A**) submitted or (**B**) published, binned into three tiers: High (JIF ≥ 42.5), Mid (JIF 15.7–27.6), and Low (JIF ≤ 9.1). Tier boundaries were defined by the actual JIFs. Within each tier, manuscripts were ordered from top to bottom first by JIF then combined *Quality* + *Impact* score. Each curve connects a manuscript’s tier position on the left to its average reviewer score on the right. Reviewer scores range from 1 (best) to 5 (worst), with lower scores indicating stronger reviews. Because *Quality* and *Impact* scores span different ranges across the dataset, each metric is displayed on its own independently scaled right-hand axis, each beginning 0.1 units below the lowest recorded score for that metric. Flatter curves indicate greater agreement between JIF tier and reviewer score. The data underlying these figures are S1_Table and https://doi.org/10.5281/zenodo.2004122.

**Fig S3.**
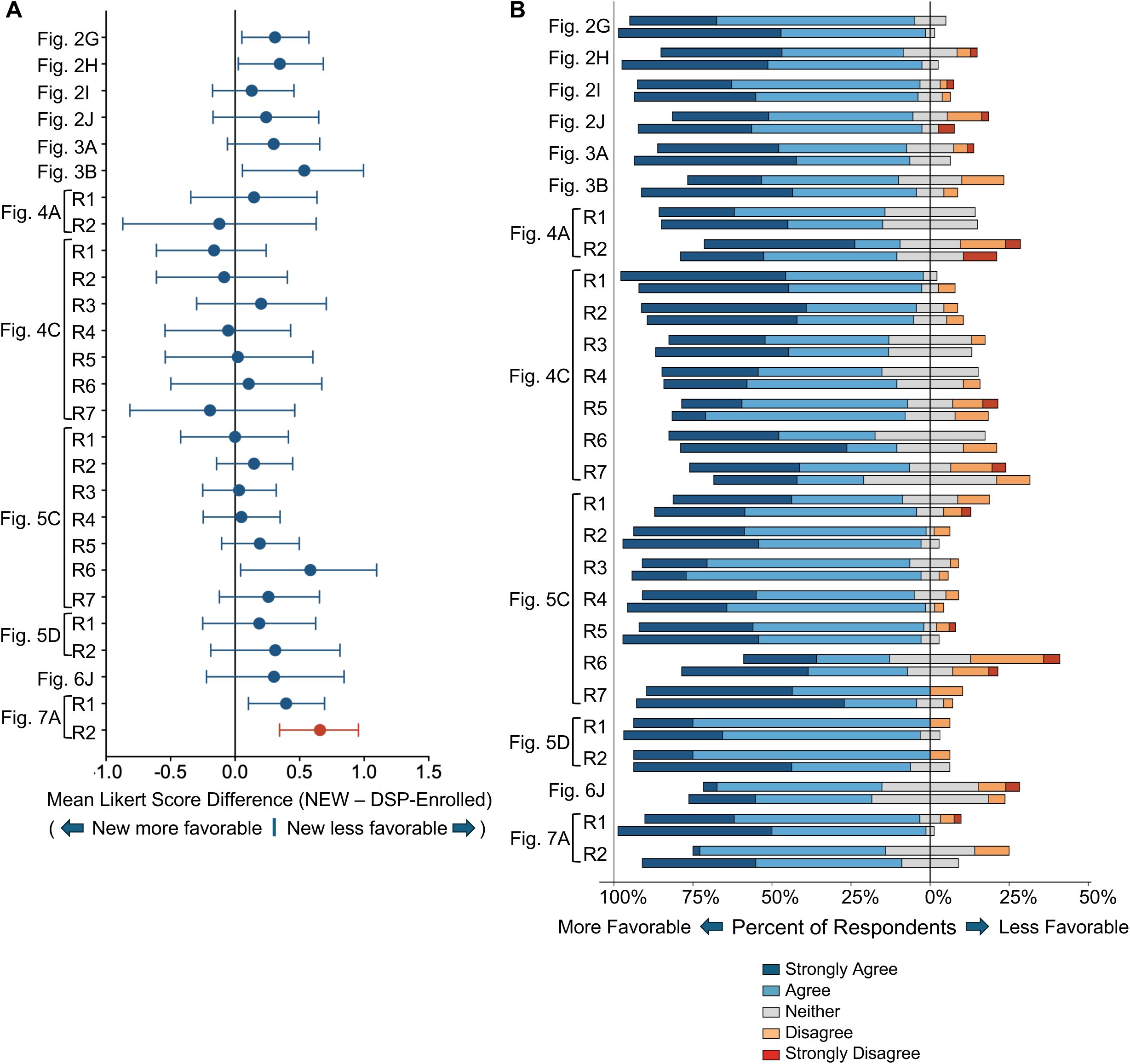
Comparison of survey responses between newly recruited (NEW) and previously enrolled (DSP-enrolled) participants. (**A**) Points represent the difference in mean Likert scores between participants new to the study (NEW) and those previously enrolled (DSP-enrolled) for each survey question. Horizontal lines indicate 95% bootstrap confidence intervals. Lower scores correspond to more favorable responses. The vertical line at zero indicates no difference between groups. (**B**) Distribution of responses for each survey question among NEW and DSP-enrolled participants. For each question, responses from NEW participants are shown in the upper bar and DSP-enrolled responses are in the lower bar. Bars are centered on the neutral response category, with more favorable responses extending to the left and less favorable responses to the right. The data underlying these figures are available in S2_Table and https://doi.org/10.5281/zenodo.2004122.

**Fig S4.**
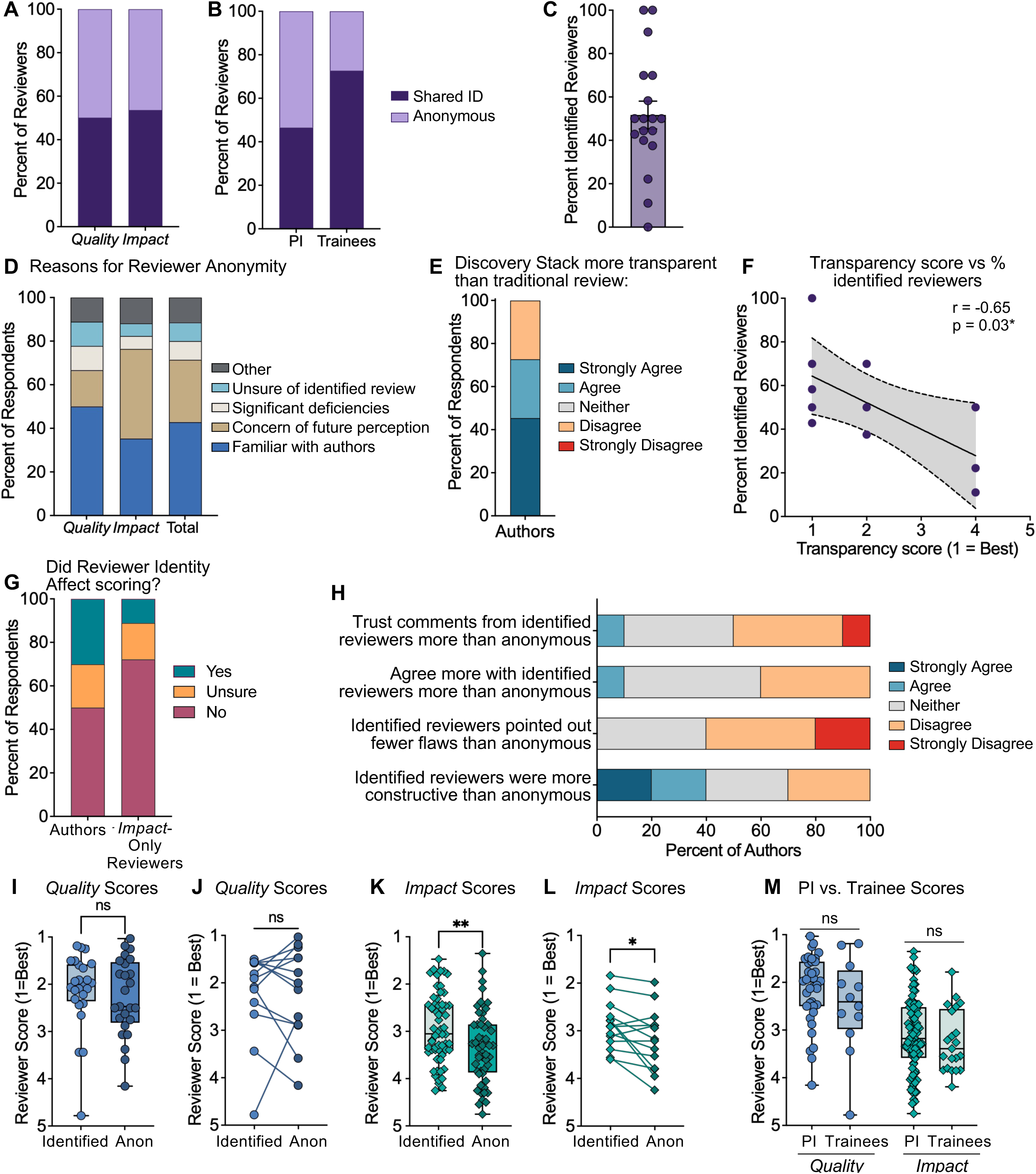
Identity Disclosure May Be Associated with Greater Perceived Transparency and Higher *Impact* Scores. (**A**) The percent of *Quality* (n=50) or *Impact*-only (n=114) reviewers who disclosed their identity or remained anonymous. (**B**) The percent of PIs (n=129) and trainees (n=33) who disclosed their identity or remained anonymous. (**C**) The percent of identified reviewers per manuscript. (**D**) Reviewers who remained anonymous (n=35) were asked a follow-up multiple-choice question regarding the reasons that influenced their choice. (**E**) Authors (n=11) were asked if they perceived the Discovery Stack review process to be more transparent than traditional review. (**F**) Transparency ratings for authors that completed surveys (n=11) were compared to percent of identified authors for each manuscript using Pearson’s correlation. (**G**) Authors (n=10) and *Impact*-only reviewers (n=18) were asked whether they noticed a difference in scoring between identified and anonymous reviewers (**H**) Authors (n=10) were asked follow-up questions regarding their perception of feedback from identified and anonymous reviewers. (**I, K**) Composite (**I**) *Quality* and (**K**) *Impact* scores were compared between identified and anonymous reviewers across all manuscripts using the Mann-Whitney test (*p*=0.0082** for *Impact*). (**J, L**) Within each manuscript, the average (**J**) *Quality* and (**L**) *Impact* scores from identified and anonymous reviewers were compared using the Wilcoxon matched-pairs signed rank test (*p*=0.026* for *Impact*). Manuscripts with no identified reviewers (*Quality* n=2, *Impact* n=1) or no anonymous reviewers (*Quality* n=3; *Impact* n=3) were excluded. (**M**) Composite *Quality* and *Impact* scores were compared between PIs (*Quality* n=37; *Impact* n=92) and trainees (*Quality* n=12; *Impact* n=21). Differences were tested using the Mann-Whitney test. The data underlying these figures are available at https://doi.org/10.5281/zenodo.2004122.

**Fig S5.**
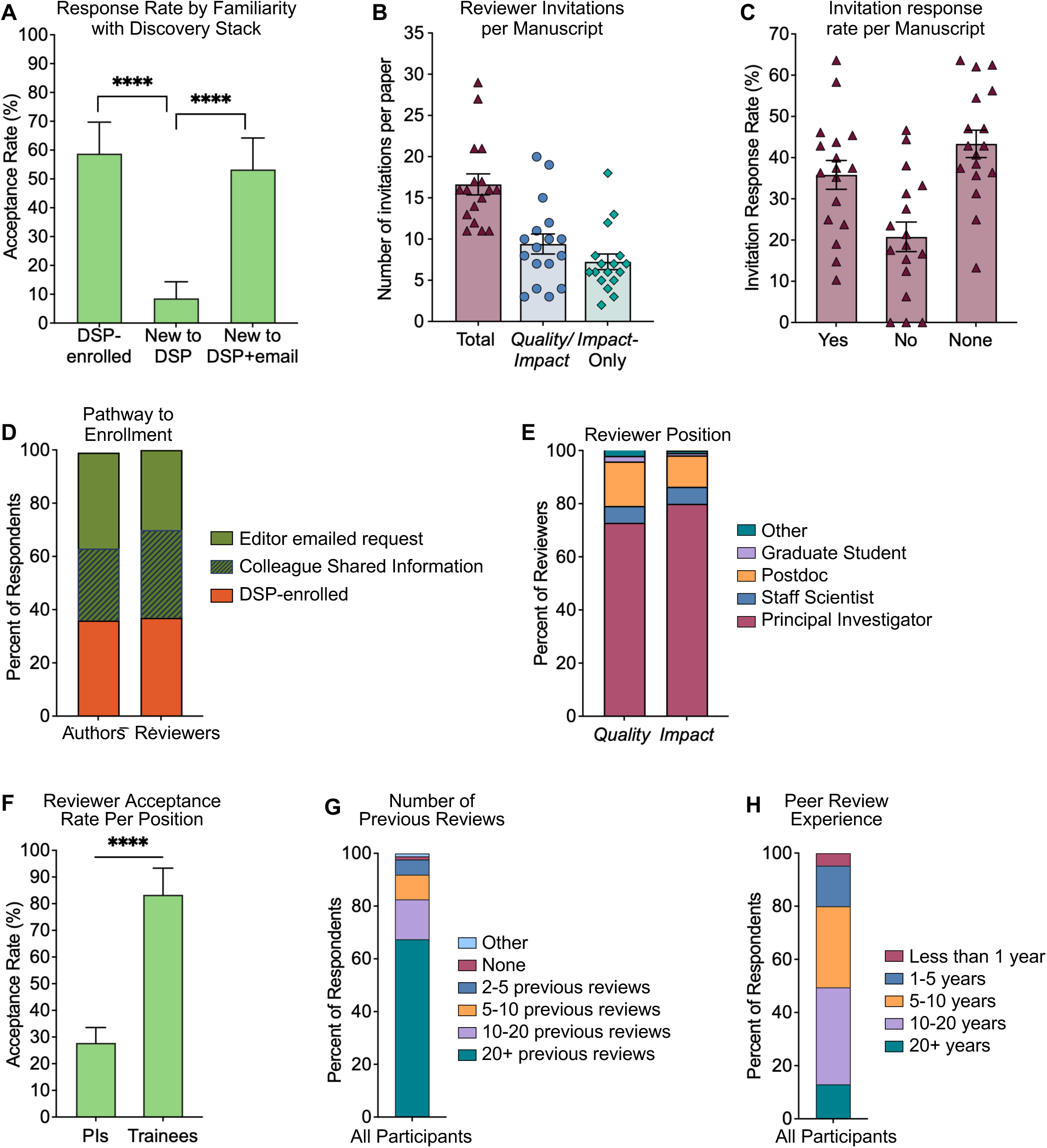
Reviewer Engagement Depends on Outreach, Networks, and Trainee Involvement. (**A**) The positive response rate was compared between individuals who were already familiar with the study (DSP-enrolled), new to the study (New to DSP), or new but contacted by a known colleague on the Scientific Advisory Board (New to DSP + email) using two-sided Fisher’s exact tests (*p*<0.0001). Odds ratios (ORs) and 95% Cis were calculated from the contingency table. P-values were adjusted for multiple pairwise comparisons using the Bonferroni correction. Error bars represent 95% CIs. DSP-enrolled vs. New to DSP (OR=15.2, 95% CI: 7.1-32.7, *p*<0.0001, adjusted), New to DSP + email vs. New to DSP vs. (OR=12.2, 95% CI: 5.8-25.7, *p*<0.0001, adjusted), DSP-enrolled vs. New to DSP + email significant (OR=1.25, 95% CI: 0.64-2.4, *p*=0.61, adjusted). (**B**) Number of reviewer invitations per manuscript to recruit *Quality/Impact* and *Impact*-only reviewers. (**C**) The percent of individuals that responded “yes”, “no”, or did not respond to reviewer invitations per manuscript. The overall acceptance was 33%. (**D**) Survey respondents indicated whether they were unfamiliar with the study, previously enrolled, or recruited by a colleague. (**E**) Percent of *Quality* (n=48) or *Impact* (n=110) reviewers at each career stage. Among *Quality* reviewers, 73% were Principal Investigators (PIs) while 80% of *Impact* reviewers were PIs. Only independent trainee reviewers (i.e., not co-reviewing with a PI) were included in these percentages. Other includes industry positions. (**F**) A multivariable logistic regression model estimated the odds of reviewer acceptance as a function of position (PI vs. trainee) and participant type (DSP-enrolled, New to DSP, New to DSP + email) as predictors. Only independent trainee reviewers, not co-reviewers, were included in the analysis. OR and 95% CI were obtained by exponentiating the logistic regression coefficients. Statistical significance was assessed using Wald tests (OR=15.3, 95% CI: 4.2-56.2, *p*<0.0001). (**G**) Survey participants reported their review experience. (**H**) The number of years of peer review experience reported by participants. The data underlying these figures are available at https://doi.org/10.5281/zenodo.2004122.

**Fig S6.**
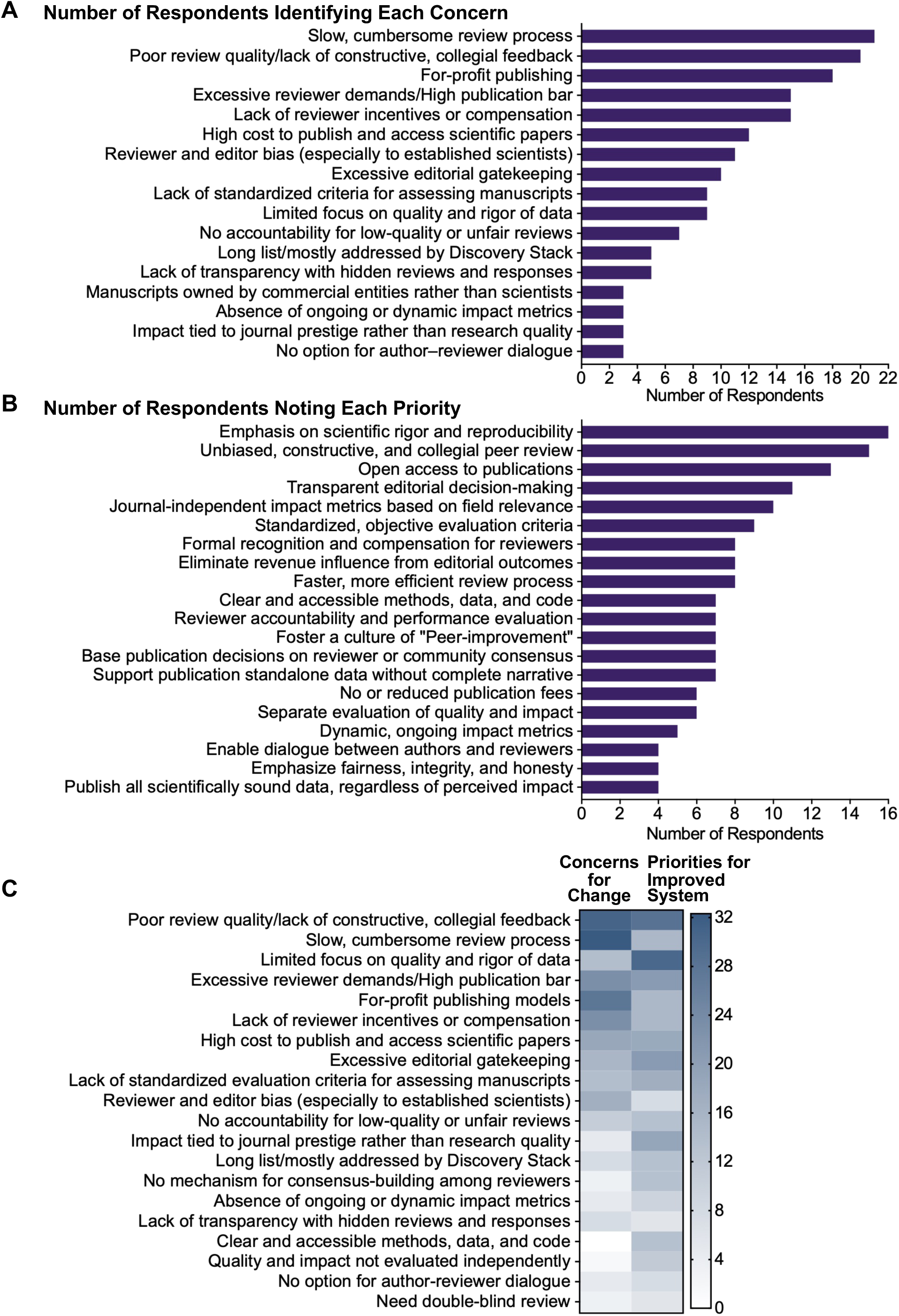
Top Priorities for Improved Scientific Publishing. (**A**) Participants (n=86) were asked what they believe needs to change about the current scientific publishing and peer review systems. A total of 65 open-ended responses were collected, categorized into key thematic areas, and tallied. The number of respondents mentioning each concern is shown. (**B**) Participants were also asked to imagine that scientific publishing didn’t exist and to identify three core principles they would prioritize if building a system from scratch. A total of 53 open-ended responses were collected, categorized by theme, and tallied. (**C**) Responses across both questions were averaged and ranked to highlight the most broadly recognized priorities for reform. The data underlying these figures are available at https://doi.org/10.5281/zenodo.2004122.

**Fig S7.**
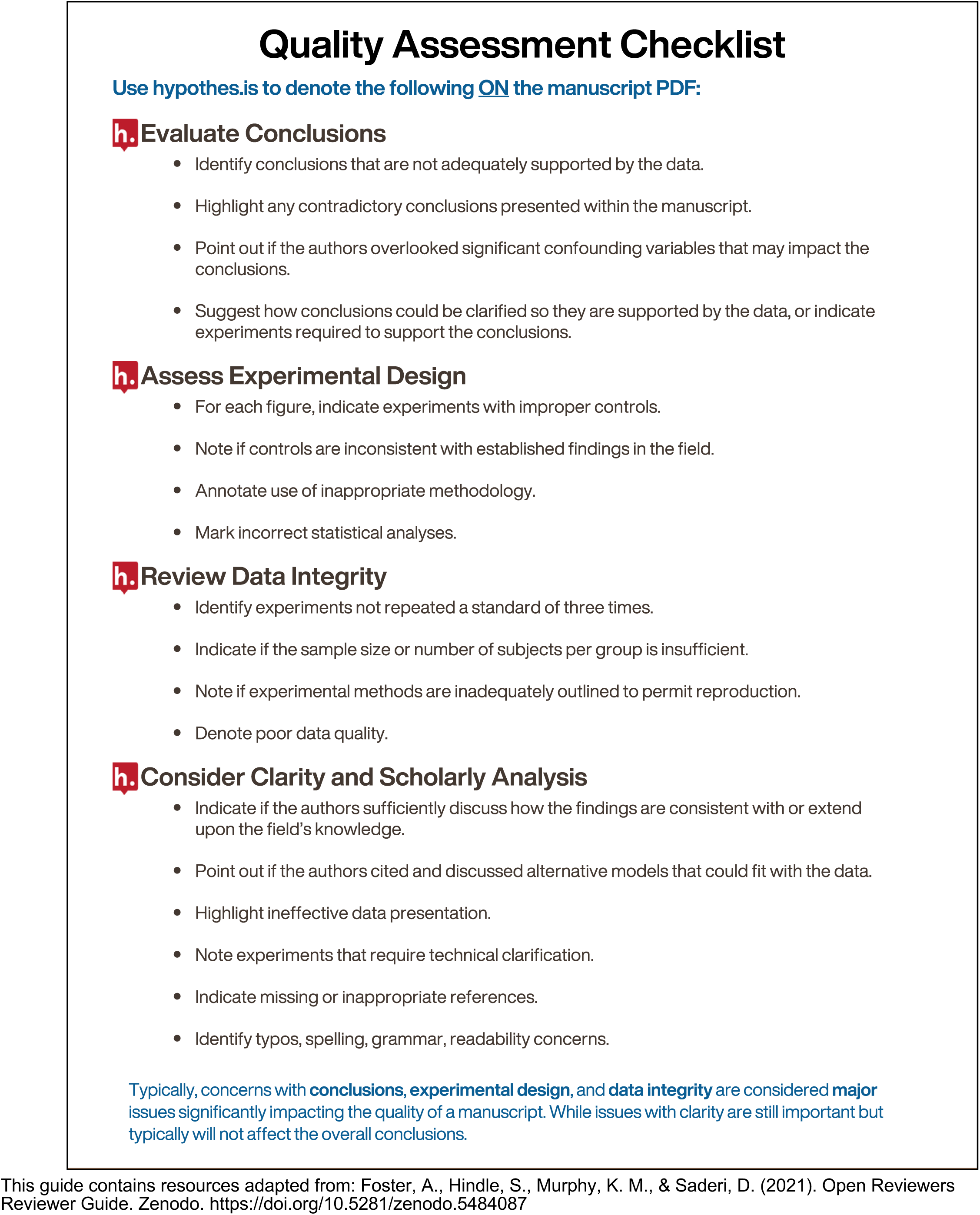
*Quality* Assessment Checklist. The *Quality* Assessment checklist was given to *Quality/Impact* reviewers to highlight key elements to consider when evaluating the *Quality* of a manuscript. It offers a comprehensive framework for reviewing conclusions, assessing experimental design, ensuring data integrity, and evaluating clarity and scholarly analysis. This guide contains resources adapted from (30) https://doi.org/10.5281/zenodo.5484087.

**S1 Table. Composite Reviewer Scores, JIFs, and Journal Review Timelines Per Manuscript.** Individual reviewer composite scores and average composite scores are shown for both *Quality* and *Impact* evaluations for each manuscript, Lower scores are more favorable evaluations. The table also shows the JIF of the journal to which each manuscript was initially submitted and ultimately published (when applicable), as well as the duration of journal review for manuscripts published in journals.

**S2 Table. Comparison of survey responses between newly recruited (NEW) and previously enrolled (DSP-enrolled) participants**. Survey responses from (NEW) and those previously enrolled (DSP-enrolled) were compared using Mann-Whitney U tests. Analyses were restricted to survey items with sufficient responses across groups. Questions that were only administered to authors were excluded due to the limited number of author respondents (NEW, n = 7; DSP-enrolled, n = 4). For all other questions, responses from all participant roles (*Quality/Impact* reviewers, *Impact*-only reviewers, authors) were pooled within the NEW and DSP-enrolled groups and compared. Lower scores corresponded to more favorable responses. P-values were adjusted using the Benjamini–Hochberg procedure. Effect sizes (r) were calculated from the standardized Mann-Whitney statistic as r = Z/√N, where Z was derived using the normal approximation with continuity and tie corrections and N is the total number of observations.

**Table S3. Bootstrap Summary Comparing Variability of *Quality* and *Impact* Reviewer Scores.** Mean differences (*Impact* - *Quality*), confidence intervals, and one-sided p-values derived from 5,000 bootstrap iterations in which *Impact* scores were randomly subsampled to match the number of *Quality* scores for each manuscript. One-sided p-values were defined as the proportion of iterations in which the mean difference (*Impact* - *Quality*) was less than or equal to zero.

